# Inhibition of cell-mediated immunity in type 1 diabetes by beta cell-targeted PD-1 agonists in pancreas tissue slices

**DOI:** 10.1101/2025.10.21.683528

**Authors:** Matthew W. Becker, Matthew Brown, Katherine Wiseman, Ana Laura Chiodetti, Mollie K. Huber, Alexandra E. Cuaycal, Pumin Sintara, Sandra M. Ferreira, Dylan Smurlick, Jessie M. Barra, Andrew M. Ladd, Denise M. Drotar, Mark A. Atkinson, Peter Weber, Hussein Al-Mossawi, Holger A. Russ, Tara M. Mahon, Todd M. Brusko, Giovanna Bossi, Edward A. Phelps

**Affiliations:** J. Crayton Pruitt Family Department of Biomedical Engineering, Herbert Wertheim College of Engineering, University of Florida, Gainesville, FL, USA; Department of Pathology, Immunology, and Laboratory Medicine, College of Medicine, University of Florida, Gainesville, FL, USA; University of Florida Diabetes Institute, University of Florida, Gainesville, FL, USA; Immunocore Ltd., Abingdon, United Kingdom; Department of Infectious Diseases and Immunology, College of Veterinary Medicine, University of Florida, Gainesville, FL, USA; Department of Pharmacology & Therapeutics, College of Medicine, University of Florida, Gainesville, FL, USA; Department of Pediatrics, College of Medicine, University of Florida, Gainesville, FL, USA; Department of Biochemistry and Molecular Biology, College of Medicine, University of Florida, Gainesville, FL, USA

## Abstract

Tissue-targeted immunotherapies for type 1 diabetes (T1D) hold potential to protect pancreatic beta cells while minimizing systemic immunosuppression. We used a bispecific agonist called Immune Modulating Monoclonal-TCR Against Autoimmune Disease (ImmTAAI), consisting of a T cell receptor (TCR) targeting domain fused with a PD-1 agonist to specifically bind beta cells and suppress autoreactive T cells. We used live pancreas slices to demonstrate targeting of ImmTAAI molecules to pre-proinsulin peptide-HLA-A2 complexes on human beta cells. ImmTAAI protected beta cells from T cell killing by increasing T cell motility and inhibiting cytokine secretion. ImmTAAI treatment also increased the motility of islet-infiltrating T cells in slices from a donor with recent-onset T1D and preserved insulin secretion in slices co-cultured with T cell avatars transduced with diabetogenic TCRs. These data demonstrate that ImmTAAI molecules have the potential to limit T cell activity locally, making this an attractive platform to elicit targeted immunoregulation in T1D.

**One Sentence Summary:** We demonstrate inhibition of cellular immunity in human type 1 diabetes using a beta cell-targeting, affinity-enhanced TCR fused to a PD-1 agonist.

## INTRODUCTION

Type 1 diabetes (T1D) is characterized by the selective destruction of insulin-producing beta cells by autoreactive T cells (*1–3*). Approaches to manipulate mechanisms of central and peripheral tolerance are actively being explored to treat T1D autoimmunity (*4–8*); however, systemic immune suppression carries significant negative drawbacks for patient health. The polyclonal autoimmune response in T1D, which at the time of diagnosis involves T and B cells recognizing multiple epitopes from an incompletely defined list of islet antigens, poses a significant challenge for the clinical success of antigen-specific immunotherapies (*9–12*). One possible solution would involve the design of therapies that promote islet-localized tolerance by targeting beta cells. Such a strategy would interface with the entire repertoire of islet-invading T cells without requisite knowledge of the autoantigen subset while avoiding the need for systemic immune suppression.

Inhibitory receptor agonists represent a class of drugs targeting peripheral immune tolerance mechanisms, which are beginning to make their way into clinical trials for autoimmune diseases (*13*). Of the inhibitory receptors, programmed death-1 (PD-1) and its ligand, PD-L1, are particularly relevant to T1D (*14*). PD-1 is expressed on infiltrating lymphocytes in non-obese diabetic (NOD) mice (*15*) and in human T1D organ donors (*16*). Disrupting PD-1/PD-L1 interactions with a monoclonal antibody leads to accelerated T1D in mice (*17–19*). Furthermore, administration of clinically approved cancer therapies that disrupt PD-1/PD-L1 interactions can lead to the development of autoimmune diabetes (*20–22*). Positive responses to drug therapies for T1D in clinical trials are also correlated with T cell exhaustion markers such as PD-1 (*23, 24*). Since PD-1 signaling in T cells depends on TCR engagement with peptide-HLA (pHLA) complexes (*25–28*), therapeutics that target tissue-specific pHLA may enable selective activation of PD-1 signaling in disease-relevant tissues, thereby promoting localized immune tolerance.

We previously designed a bispecific PD-1 agonist Immune modulating monoclonal TCR Against Autoimmunity (ImmTAAI) molecule with a targeting end composed of a soluble TCR that is highly specific for a pre-proinsulin (PPI) peptide, PPI_15-24_, bound to HLA-A*02:01 (HLA-A2) on beta cells, together with an effector end composed of an agonistic PD-1 binding nanobody (*29*). PD-1 agonist ImmTAAI molecules are potent inhibitors of effector T cells and reduce cytokine secretion and beta cell killing *in vitro* (*29*). T cell inhibition only occurs when PD-1 agonist ImmTAAI molecules are bound to target cells, an activity that does not occur when ImmTAAI molecules are in solution (*29*). However, these studies were performed with a homogeneous beta cell line; hence, further investigation is needed to understand the scope of bispecific ImmTAAI molecules in complex primary human tissues. One novel tool to study ImmTAAI molecules is that of pancreatic tissue slices. These explants are thin slices (∼120 μm thickness) of live tissue from transplant-quality donor organs, embedded in agarose and maintained in a functional state in vitro (*30, 31*). Each tissue slice contains both exocrine and endocrine tissue with intact islets capable of glucose sensing and insulin secretion (*32, 33*). Slices also preserve islets in the native tissue context in disease states such as T1D where the islet may be infiltrated and structurally compromised (*33–35*).

In this study, we investigated whether ImmTAAI molecules maintain beta cell and HLA-A2 specificity within the heterogeneous tissue of pancreas slices. Through immunofluorescent labeling and confocal microscopy of live tissue, we observed that ImmTAAI molecules maintain specificity with minimal to no off-target binding. ImmTAAI molecules also protected beta cells from killing by T cell avatars transduced with diabetogenic TCRs. ImmTAAI demonstrated effective binding to beta cells in a recent-onset T1D donor, resulting in changes to T cell motility and reducing T cell:beta cell interactions, indicating that this therapy may represent a viable strategy for reversing autoimmunity.

## RESULTS

### Fluorescently labeled ImmTAAI molecules bind target pHLA and inhibit TCR signaling

We generated cyanine-based far-red fluorescent dye (CF647) labeled ImmTAAI molecules specific for PPI peptide PPI_15-24_ pHLA-A2 and for telomerase (Tel) peptide hTERT_540–548_ pHLA-A2 (negative control) to visualize binding in live pancreas slices. We conducted Surface Plasmon Resonance (SPR) studies to determine binding kinetics for labeled and unlabeled PPI ImmTAAI and Tel ImmTAAI to their corresponding surface-immobilized cognate pHLA. We obtained single cycle kinetics profiles, where a series of five ImmTAAI concentrations were sequentially injected over immobilized pHLA, followed by a long dissociation phase (Fig. 1A). Both labeled and unlabeled PPI ImmTAAI demonstrated similar affinity for cognate pHLA in the picomolar range, with the time taken for half the bound ImmTAAI to dissociate from pHLA (t_1/2_) > 6 hours (Fig. 1A). Similarly, both labeled and unlabeled Tel ImmTAAI retained picomolar affinity for cognate pHLA with t_1/2_ > 4 hours (Fig. 1A). Hence, ImmTAAI molecules strongly bind their cognate pHLA, and pHLA binding was not significantly affected by the CF647 fluorescent tag.

**Fig. 1.**
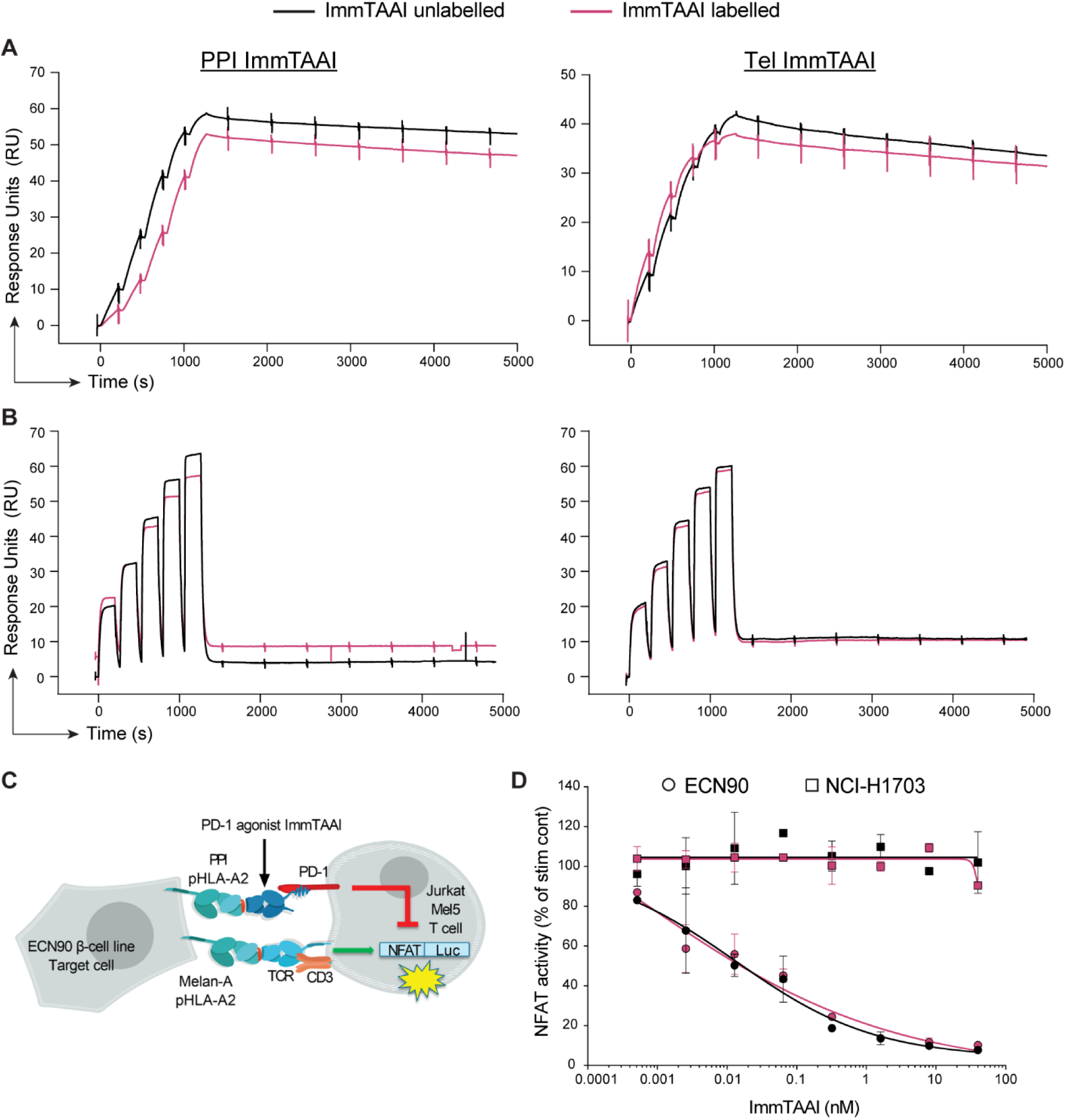
Fluorescently labeled ImmTAAI molecules bind target pHLA and inhibit TCR signaling. (**A**) CF647-labeled ImmTAAI molecules were assessed for pHLA binding and compared to unlabeled molecules by SPR using BIAcore 8K. Binding to PPI or Tel cognate pHLA was carried out at 37°C. (**B**) Similarly, CF647-labeled ImmTAAI molecules were assessed for PD-1 binding and compared to unlabeled molecules. PD-1 binding was carried out at 25°C. (**C**) Schematic of the ECN90 beta cell line: Jurkat NFL Mel5 PD-1 reporter assay. (**D**) ECN90 or NCI-H1703 cells were pulsed with Melan-A activating peptide, and titrations of unlabeled or labeled PPI ImmTAAI molecules were added.

Next, we assessed the effect of CF647 labeling on ImmTAAI effector region binding to PD-1. We obtained single cycle kinetics profiles where a series of five PD-1 concentrations were sequentially injected over immobilized ImmTAAI. The anti-PD-1 portion of the ImmTAAI molecule displayed nanomolar affinity with a relatively short t_1/2_ of ∼13 seconds (Fig. 1B). Binding to PD-1 was likewise unaffected by labeling with CF647 for both PPI and Tel ImmTAAI molecules (Fig. 1B).

To identify any potential impact of CF647 labeling on cellular potency or selectivity, the labeled and unlabeled PPI ImmTAAI molecules were also tested in a Jurkat NFL Mel5 PD-1 reporter assay (Fig. 1C) (*29*). This incorporated the ECN90 human beta cell line as targets with natural PPI pHLA-A2 presentation and NCI-H1703 human squamous cells that do not present the PPI epitope. Target cells were pulsed with the Melan-A antigen to engage the Mel5 TCR on the Jurkat cells and evaluate the inhibitory effect of PD-1 engagement by the targeted ImmTAAI on T cell activity. Both labeled and unlabeled PPI ImmTAAI molecules inhibited cellular TCR signaling with comparable potency (Fig. 1D). No inhibitory activity at ImmTAAI concentrations up to 40 nM was observed when using NCI-H1703 target cells (Fig. 1D). Overall, these data demonstrate that the CF647 labeling did not impact ImmTAAI potency or selectivity.

### ImmTAAI binding and labelling of beta cells in pancreas slices is concentration-dependent

Live tissue slices of 120 µm thickness were generated by the Network for Pancreatic Organ donors with Diabetes (nPOD) program (*36, 37*) from human pancreas organ donors using a vibratome. Donor information is listed in Table S1. Pancreas slices contained islets in the native tissue microenvironment as visualized by darkfield stereomicroscopy (Fig. 2A). Pancreas slices from a control donor without diabetes expressing HLA-A2 were incubated with CF647-labeled PPI ImmTAAI, fixed, stained for insulin, glucagon, and the beta cell surface marker ectonucleoside triphosphate diphosphohydrolase-3 (ENTPD3) (*34, 38–40*) (Fig. 2B). ImmTAAI colocalized with ENTPD3 in the plasma membrane of beta cells. Next, we incubated live slices from an HLA-A2-positive donor without diabetes with CF647-labeled ImmTAAI molecules at different concentrations for 1 hour and imaged the slices by live cell confocal microscopy (Fig. 2C, S1). Islets in live slices were located using a combination of reflected light backscattered off the insulin granules (*41*) and co-staining for ENTPD3. The control Tel ImmTAAI did not label the tissue. Incubation with 2, 20, and 200 nM PPI ImmTAAI resulted in islet-specific labeling with a dose-dependent increase in mean fluorescence intensity (MFI) (Fig. 2D). PPI ImmTAAI co-localized with ENTPD3 starting at 2 nM, whereas the weak fluorescent signal detected from Tel ImmTAAI or unlabeled slices did not co-localize with ENTPD3 (Fig. 2E). Image quantification for MFI showed that 20 nM ImmTAAI was required to achieve significant increases in islet-localized ImmTAAI brightness relative to control or unlabeled samples due to frequent autofluorescent background spots in the islet tissue, which we attribute to accumulated lipofuscin in the long-lived post-mitotic beta cells (*42*). Incubation with ImmTAAI molecules did not negatively affect cell viability in slices (Fig. S1). Taken together, these data indicate that PPI ImmTAAI binds specifically to beta cell surfaces in a concentration-dependent manner in the complex environment of live native pancreatic tissue.

**Fig. 2.**
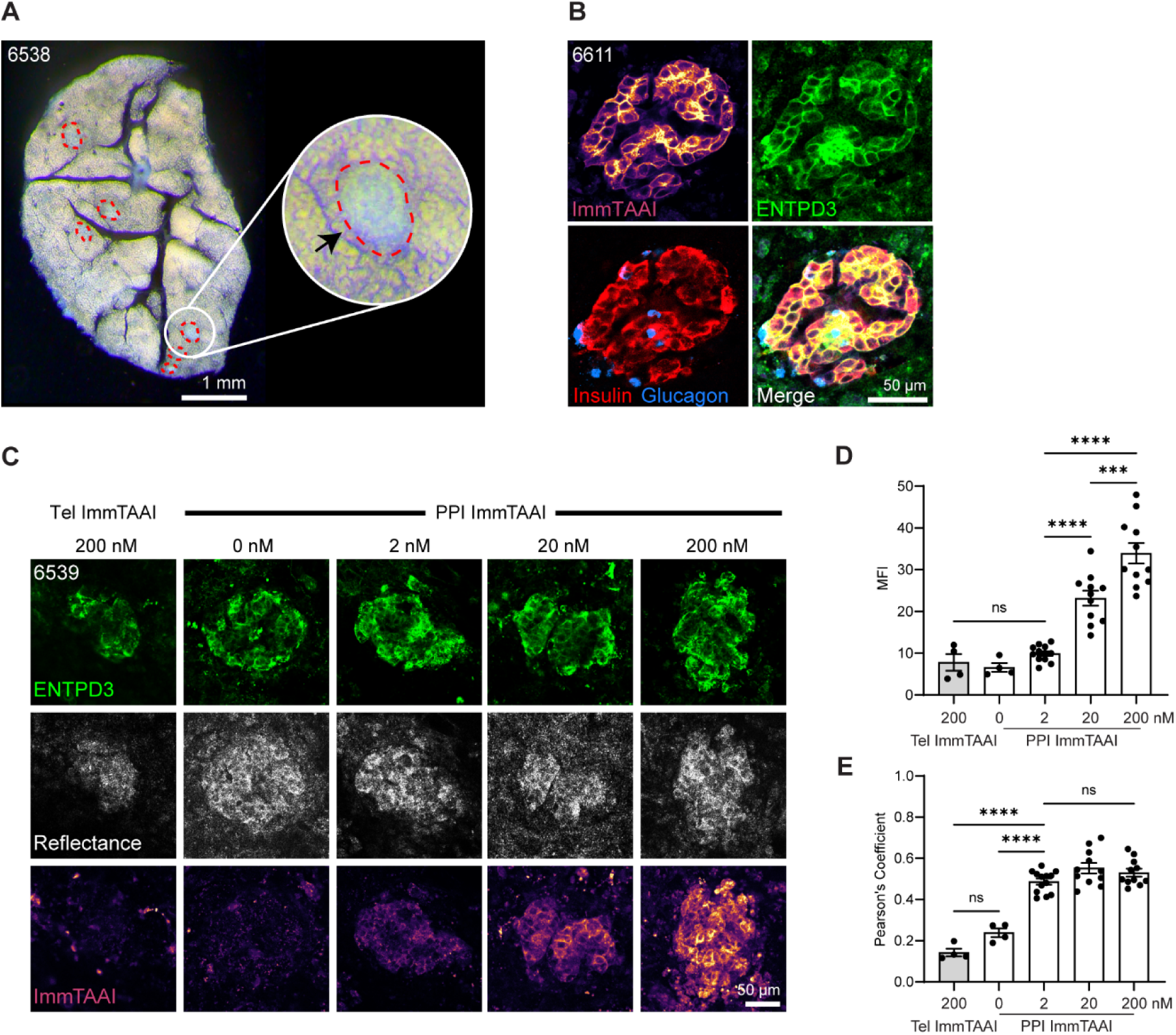
ImmTAAI binding and labelling of beta cells in pancreas slices is concentration-dependent. (**A**) Darkfield stereomicroscopy image of a pancreas slice showing islets outlined in red. (**B**) Pancreas slice incubated with CF647 labeled PPI ImmTAAI and subsequently fixed and stained for insulin, glucagon, and ENTPD3, and imaged by confocal. (**C**) Confocal microscopy images of live pancreas slices incubated with CF647-labeled PPI ImmTAAI at 2, 20, or 200 nM for 1 hour and stained with a labeled primary antibody for cell surface ENTPD3. The off target CF647 labeled Tel ImmTAAI was used as a control. (**D**) Mean fluorescent intensity (MFI) quantification of confocal microscopy images showing the dose-dependent binding of PPI ImmTAAI to tissue slices. (**E**) Pearson’s Coefficient co-localization analysis of ImmTAAI fluorescent signal with anti-ENTPD3 fluorescent signal. PPI ImmTAAI co-localizes with anti-ENTPD3 well at all three concentrations tested, indicating beta cell specific binding is maintained in native tissue. Individual dots represent measurements per islet (means from 1 donor for Tel ImmTAAI, 2 donors for PPI ImmTAAI labeled). Statistical differences were determined by two-way ANOVA followed by Tukey’s post-hoc analysis. ns = not significant, *** p < 0.001, **** p < 0.0001.

### ImmTAAI binding is HLA-specific in tissue slices

The TCR domain of ImmTAAI molecules specifically targets PPI_15-24_ presented in the context of HLA-A2. To confirm the HLA specificity of the ImmTAAI in primary tissue, we compared its binding in pancreatic slices from an HLA-A2-positive donor and an HLA-A2-negative donor (HLA-A*31). Tissue slices were incubated with 20 nM CF647-labeled PPI or Tel ImmTAAI for two hours and labeled with anti-ENTPD3 before live imaging (Fig. 3A). There was no specific fluorescent signal in tissue from the HLA-A*31 donor for either PPI or Tel ImmTAAI, while significant binding occurred with PPI ImmTAAI in the HLA-A2 donor (Fig. 3B). We again confirmed that this binding was beta cell-specific by quantifying co-localization with AF594-labeled anti-ENTPD3 (Fig. 3C).

**Fig. 3.**
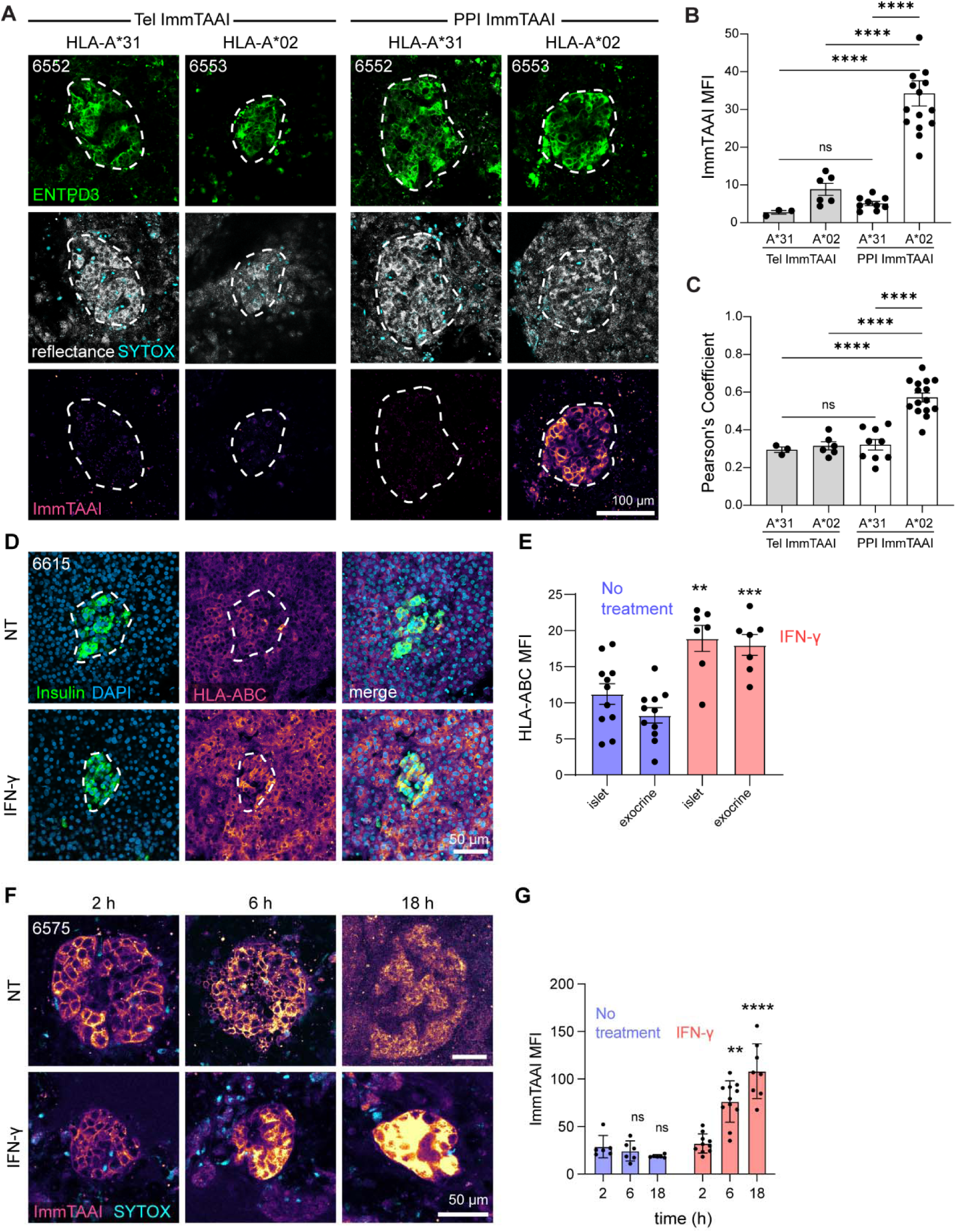
ImmTAAI binding is HLA-specific in tissue slices. (**A**) Confocal microscopy images of pancreas slices incubated with 20 nM CF647-labeled PPI ImmTAAI for 2 hours and co-stained for ENTPD3. No ImmTAAI staining is observed in slices from HLA-A*31 tissue. Islet granularity (reflectance) and viability (SYTOX blue) are also shown. (**B**) MFI analysis of ImmTAAI binding in HLA-A2 negative and positive tissue, showing binding only in HLA-A2 positive tissue. (**C**) Pearson’s Coefficient co-localization analysis of ImmTAAI fluorescent signal with anti-ENTPD3 fluorescent signal. ImmTAAI signal co-localizes with ENTPD3 only in HLA-A2 positive tissue. Each dot represents a separate islet from 1 or 2 separate slices per condition (n = 1 donor for Tel ImmTAAI in HLA-A*31 tissue, n = 2 donors for HLA-A2) and is the mean of 3-6 z-planes taken per islet. (**D**) Confocal microscopy images of pancreas slices treated with a 5-hour pulse of 100 IU/ml IFN-γ, then fixed after 24 hours and stained for insulin and HLA-ABC. (**E**) MFI analysis of HLA-ABC expression in islet and exocrine regions. Each dot represents a separate islet / exocrine region image from ≥3 slices per condition. (**F**) Live confocal microscopy images of pancreas slices treated with a 5-hour pulse of 100 IU/ml IFN-γ and incubated with CF647-labeled PPI ImmTAAI at 2, 6, or 18 hours. (**G**) MFI analysis of ImmTAAI signal at each timepoint post IFN-γ treatment. Each dot represents a separate islet from ≥3 slices per condition. NT = no treatment. Statistical differences were determined by two-way ANOVA followed by Tukey’s post-hoc analysis. ns = not significant, **** p < 0.0001.

T1D is characterized by HLA class I hyperexpression within islets (*43, 44*). HLA class I expression can be induced in pancreatic slice cultures by treatment with exogenous interferon gamma (IFN-γ) (*45*). Thus, we pre-treated slices with a pulse of 100 U/ml of IFN-γ for 5 hours, washed off the cytokine, and assessed islets after 18 hours for HLA class I upregulation. Immunostaining of fixed slices confirmed a robust upregulation of HLA class I in both islet and exocrine tissues (Fig. 3D,E). ImmTAAI binding accumulated over time following IFN-γ treatment (Fig. 3F,G). Together, these results show that ImmTAAI binding to beta cell surfaces depends on cell surface expression of pHLA, and IFN-γ-mediated inflammation results in increased ligation of ImmTAAI on the surface of beta cells.

### PPI ImmTAAI suppresses T cell engagement with beta cells to prevent cell-mediated killing

Antigen recognition by T cells results in swarming behavior and T cell arrest on target cells in a manner dependent on TCR engagement (*46*). The PD-1/PD-L1 pathway has been implicated in promoting immunological tolerance by suppressing the TCR-driven stop signal (*18, 19*). PD-1/PD-L1 complexation reduces the interaction time and lowers cytokine production when CD4^+^ T cells engage with antigen presenting cells (APCs) (*47*). Conversely, blockade of PD-1/PD-L1 interactions decreases T cell migration velocity, enhances cytokine production, and boosts TCR signaling and activation (*19*). Therefore, PD-1 agonist ImmTAAI bound to beta cells could increase the mobility of T cells and protect beta cells from autoreactive T cell-mediated killing. To measure this effect, migration velocity was tracked in PPI_6-14_-specific T cells co-cultured with EndoC-βH2 beta cells. The PPI_15-24_ ImmTAAI targeted to beta cells enhanced the mobility of T cells by reducing the time of interaction between T cells and target cells (Fig. 4 A,B) while the untargeted Tel ImmTAAI molecule did not affect T cell speed as compared to the no ImmTAAI molecule condition (Fig. 4 A,B). To determine whether there is a correlation between T cell mobility and beta cell killing, the beta cell viability was assessed over time (Fig. 4 C,D). As previously reported (*29*), PPI ImmTAAI suppressed T cell killing and preserved beta cell growth *in vitro* (Fig. 4 C,D). Beta cell protection was not detected with untargeted Tel ImmTAAI or in the absence of ImmTAAI (Fig. 4 C,D). Taken together, the PPI ImmTAAI suppressed T cell function and enhanced T cell mobility preventing beta cell destruction.

**Fig. 4.**
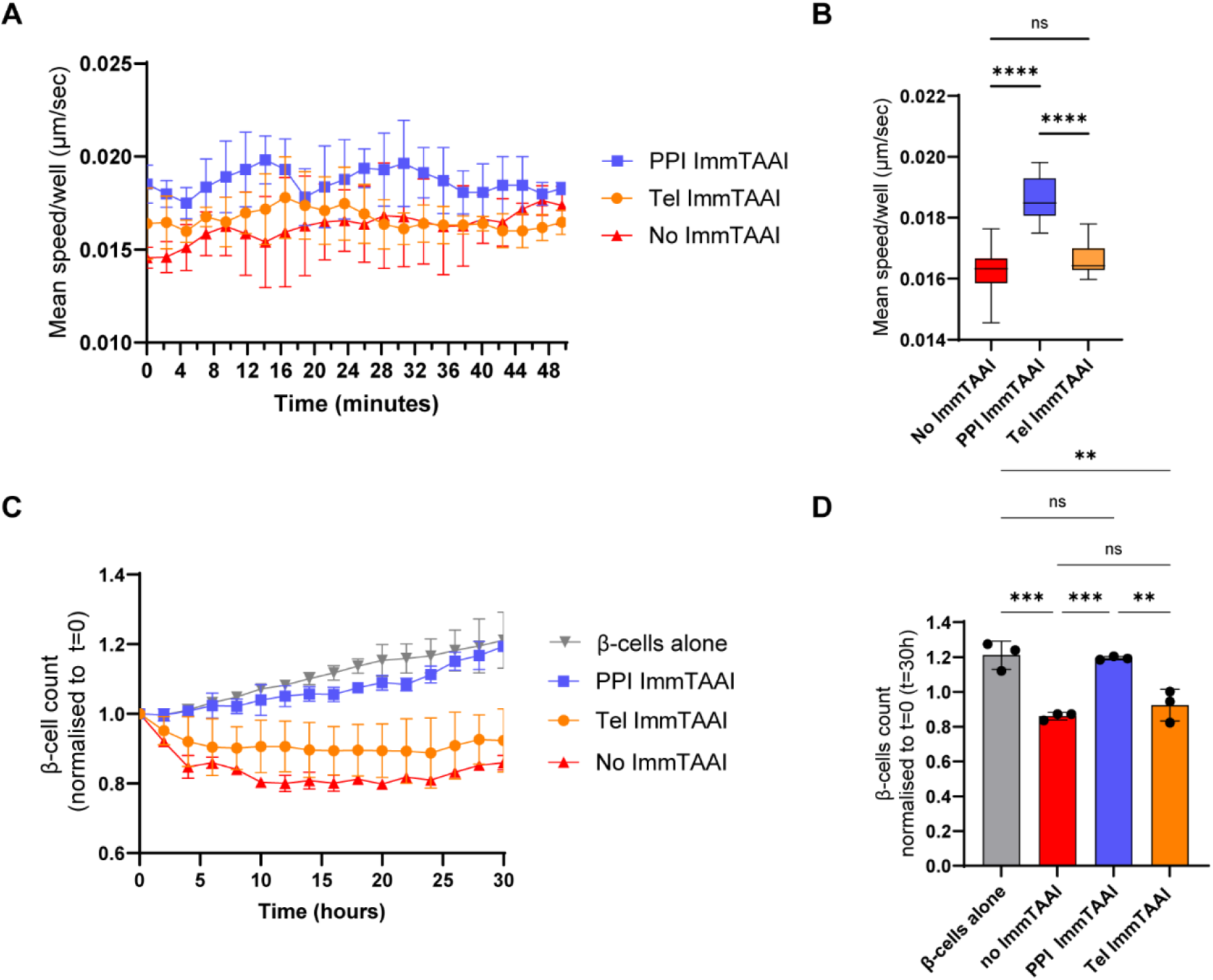
PPI ImmTAAI suppresses T cell engagement with beta cells to prevent cell-mediated killing. **(A**) Time series line plot representing the mean speed/well of T cell clone 4b (PPI_6-14_ specific) in the presence of EndoC-βH2 red cells, 10 nM PPI ImmTAAI, 10 nM Tel ImmTAAI, or without molecule. (**B**) Boxplot representing the mean speed/well of all the timepoints shown in panel A (ns: not significant, **** p<0.0001; one-way ANOVA with multiple comparisons analysed by Tukey’s test). Data are representative of three independent experiments. (**C**) Growth curves of EndoC-βH2 red cells in the presence of T cell clone 4b at 1:1 E to T ratio, 10 nM PPI ImmTAAI, 10 nM Tel ImmTAAI or no molecule over time. Time series line plot of live EndoC-βH2 red cell number over time normalised to the initial cell count at time=0. (**D**) Bar plots representing the mean/well ± SD of live EndoC-βH2 red cells count at time = 30 hours normalised to the initial cell count at time = 0. Statistical differences were determined by one way ANOVA with multiple comparisons analysed by Tukey’s test. ns = not significant, *** p<0.001, **p<0.01; Data is representative of three independent experiments.

### PPI ImmTAAI reduces T cell stopping in live pancreas slices from a recent-onset T1D donor

To confirm that the PPI ImmTAAI maintains its beta cell binding capacity in donors with recently diagnosed T1D, pancreas slices from an HLA-A2-positive donor with recently diagnosed T1D (nPOD 6551, 20 years of age, 0.58 years T1D duration, Table S1) and remaining beta cells were labeled with 20 nM CF647-labeled PPI ImmTAAI for two hours and co-stained for ENTPD3. We observed robust ImmTAAI binding to islets in slices from this donor (Fig. S2). Although nPOD donor 6551 was reported to have insulitis in some islets (nPOD histopathology database and (*34*)), we were unable to record live slices with insulitis and ImmTAAI labeling in this donor, consistent with the variable intensity and frequency of insulitis that is well-documented in human T1D (*3, 48*). We identified a second pancreas slice donor with new-onset T1D (nPOD 6578, 11 years of age, 0 years T1D duration), whose islets had both high levels of T cell infiltration and remaining beta cells (Fig. 5A). We treated slices with 20 nM CF647-labeled PPI ImmTAAI overnight, stained endogenous T cells with an antibody for CD3, and beta cells with an antibody for ENTPD3, then measured the effect of the ImmTAAI on CD3^+^ T cell infiltrated islets. We generated 30-minute timelapse recordings by confocal microscopy to track T cell motility (Fig. 5B). In slices not treated with ImmTAAI, we observed many T cells with low motility within the islet boundary, while T cells outside of the islet had high motility (Fig. 5C,D). The behavior of the slow-moving T cells, which remained within a confined localization, is suggestive of specific TCR-HLA Class I interactions promoting T cell stopping, consistent with our previously reported observations in T1D slice donors (*34*). In slices treated with PPI ImmTAAI, the mean T cell speed within islets matched that of the acinar tissue (Fig. 5C,D). These data suggest that PD-1 ligation by beta cell surface bound ImmTAAI molecules increases the motility of the endogenous T cell infiltrate in islets of a donor with recent-onset T1D.

**Fig. 5.**
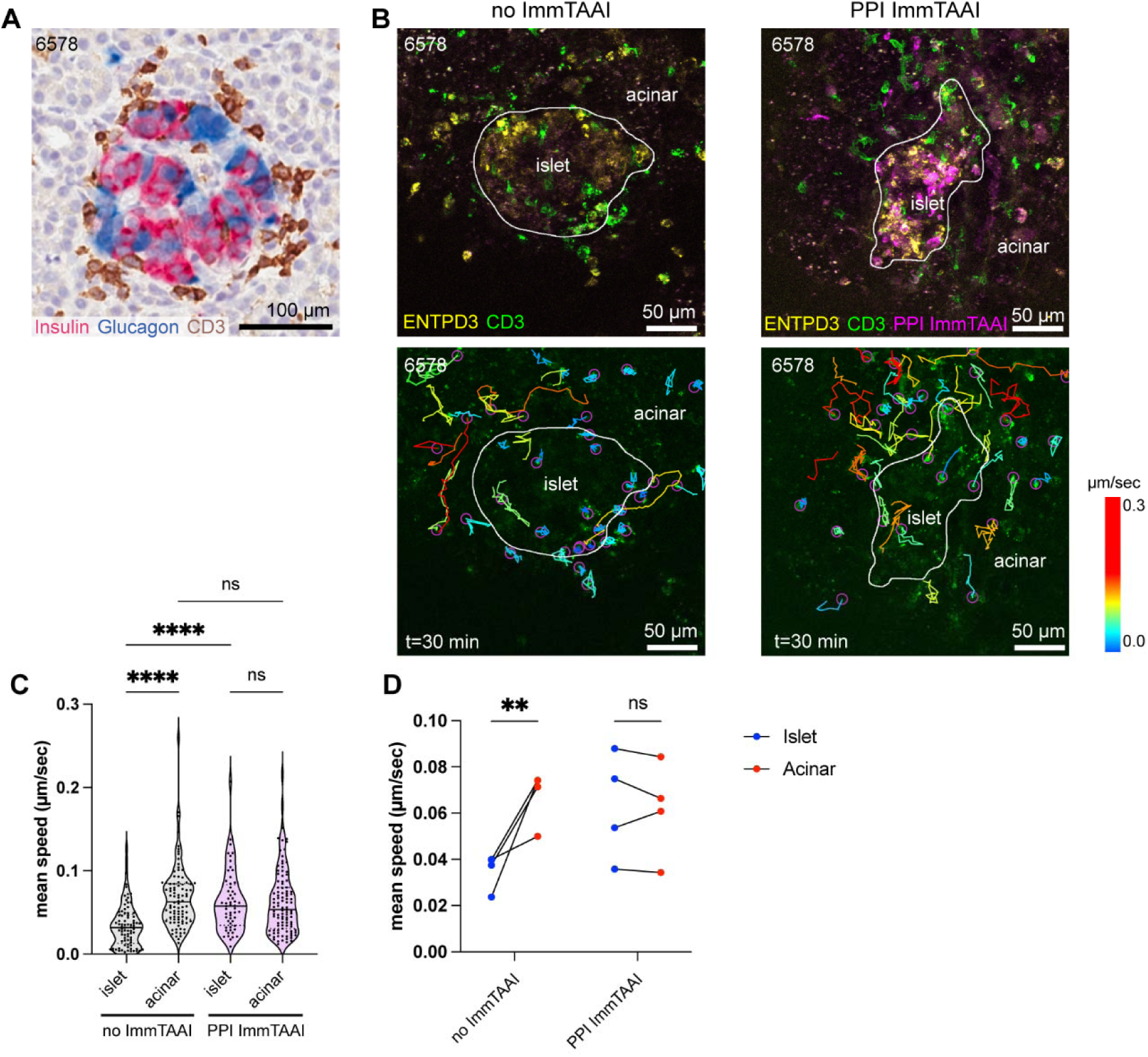
PPI ImmTAAI reduces T cell stopping in live pancreas slices from a recent-onset T1D donor. (**A**) Representative immunohistochemistry image from the nPOD histopathology database (https://portal.jdrfnpod.org/) of an insulitic islet from at-onset T1D donor 6578. (**B**) Still frames from live cell confocal microscopy videos depicting ENTDP3+ islets with CD3+ infiltrates with and without ImmTAAI (top panels) and corresponding T cell tracks over 30 minutes. (**C**) Quantification mean speed of total T cell tracks inside and outside of the islet boundary (white outline in panel B) pooled from 4 islet recordings each with and without ImmTAAI treatment.; two-way ANOVA with multiple comparisons analysed by Tukey’s test. (**D**) Average T cell speed per islet recording inside and outside of the islet boundary with and without ImmTAAI treatment. Statistical differences were determined by two-way ANOVA with multiple comparisons analysed by Turkey’s test. ns = not significant, **** p<0.0001, ** p<0.01.

### PPI ImmTAAI suppresses killing of stem cell-derived beta cells by T cell avatars

To understand the ability of ImmTAAI molecules to functionally inhibit effector T cells, we implemented a model of T cell mediated killing. This system consisted of antigen-specific T cell “avatars,” generated via lentiviral TCR gene transfer to primary human CD8^+^ T cells (*49–51*) and human stem cell-derived beta cells (sBCs) differentiated from an HLA-A2-positive line to serve as target cells (Fig. 6A) (*52, 53*). We generated two different T cell avatars specific to beta cell antigens islet-specific glucose-6-phosphatase catalytic subunit related protein (IGRP_265-273_) and PPI_6-14_. These TCRs target different peptides than ImmTAAI molecules (PPI_15-24_), ensuring that downstream functional assays specifically test PD-1 engagement rather than simply masking TCR-pHLA interactions. We also generated a negative control avatar to determine the background activity of the T cell avatars using a TCR specific for melanoma antigen (MelanA_27-35_ also known as MART-1) (*54*), an antigen not expressed in beta cells. Furthermore, we generated a positive control avatar expressing an HLA-A2-specific chimeric antigen receptor (HLA-A2-CAR) that directly recognizes HLA class I on target cells (*55*) to determine the maximal activity of the avatars.

**Fig. 6.**
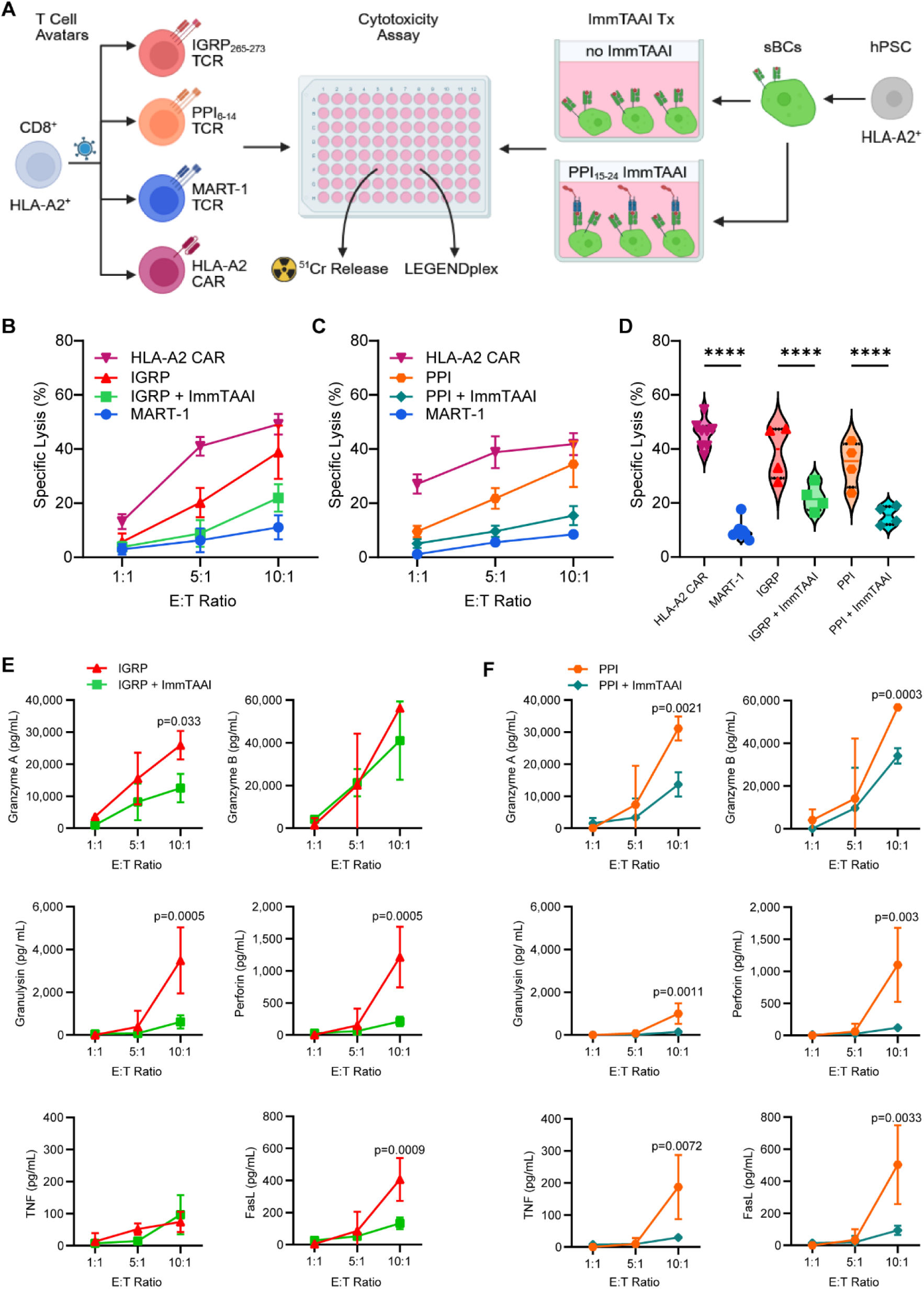
PPI ImmTAAI suppressed killing of stem cell derived beta cells by T cell avatars. (**A**) Experimental scheme depicts the co-culture assay system used to evaluate the immunogenicity of sBC treated with ImmTAAI in the presence of HLA-A2 CAR, MelanA_27-35_-reactive (MART-1), IGRP_265-273_-reactive (Clone 32), or PPI_6-14_-reactive (clone 15b) CD8^+^ avatars. (**B**) Specific lysis of sBCs by IGRP avatars after 24 hours of co-culture or (**C**) PPI avatars after 48 hours of co-culture with or without ImmTAAI treatment at each effector:target (E:T) ratio. n=4. (**D**) Specific lysis for the 10:1 E:T ratio compared across all groups. **** p< 0.0001 by two-way ANOVA with multiple comparisons. n=4. (**E,F**) Line plots show cell culture supernatant concentrations of pro-inflammatory T cell cytokines (granzyme A, granzyme B, granulysin, perforin, TNF, FasL) produced by T cell avatars. P-values are reported for paired samples using two-way ANOVA with Bonferroni’s correction for multiple comparisons.

We validated the activation of T cell avatars by kinetic analysis of CD69 and PD-1 upregulation following stimulation with anti-CD3/anti-CD28 Dynabeads (Fig. S3). We next verified that PPI ImmTAAI natively bound to the sBCs without need for exogenous peptide loading (Fig. S4). We then co-cultured the T cell avatars and sBCs at varying effector to target ratios in the presence or absence of 20 nM PPI ImmTAAI. We performed a chromium release assay to determine the impact of PPI ImmTAAI on the cytotoxicity of the T cell avatars (Fig. 6B,C). To perform this assay, target beta cells were loaded with radioactive chromium-51 (^51^Cr). When the target cells are lysed by immune effectors the ^51^Cr is released. The amount of ^51^Cr in the media is compared to a spontaneous release control (target cells alone) to determine the cytotoxic activity of the effector cells (% specific lysis). Specific lysis of sBCs by IGRP- and PPI-specific avatars correlated with the number of T cell avatars and was similar to the HLA-A2-CAR positive control at the highest effector to target ratio, while T cells specific to the negative control antigen MART-1 demonstrated significantly less sBC killing (Fig. 6B-D). The small number of sBCs destroyed by MART-1 specific T cells is most likely attributable to allorecognition by the endogenous TCR still present in the T cell avatars. Importantly, ImmTAAI molecules significantly reduced specific lysis by both IGRP- and PPI-specific T cell avatars (Fig. 6B-D).

Multiplex cytokine assays were performed on the supernatants from the T cell avatar-sBC co-culture experiments (*56*). PPI ImmTAAI treatment significantly downregulated the expression of an array of effector molecules produced by cytotoxic T cells, including granzyme A, granulysin, perforin, and FasL for IGRP-specific avatars (Fig. 6E) and granzyme A, granzyme B, granulysin, perforin, TNF, and FasL for PPI-specific avatars (Fig. 6F). Together, these results demonstrate that ImmTAAI effectively protects against beta cell killing by CD8^+^ T cells expressing islet autoreactive TCRs.

### PPI ImmTAAI suppresses killing of primary islets in human pancreas slices by T cell avatars

Human pancreas slices with T cell infiltration of islets are rare specimens, and we were unable to clearly observe beta cell killing by endogenous immune cells in human pancreas slices from recent-onset T1D donors. We speculate this is because the process of beta cell loss in human T1D occurs over a prolonged period of months to years (*57*), while we are only able to maintain pancreas slices in culture for several days. To further evaluate ImmTAAI molecules’ ability to protect beta cells from T cell killing in the human pancreas tissue, we adapted the *in vitro* cell-mediated killing assay (*49, 50, 58*) (Fig. 6A) into a new *ex vivo* model of T1D by co-culturing T cell avatars specific for beta cell antigens (IGRP_265-273_ and PPI_6-14_) with pancreas slices from HLA-A2-positive donors without diabetes (Fig. 7A, Table S1). Slices were incubated with 200,000 T cells per slice, with or without 20 nM PPI ImmTAAI, then evaluated by live imaging for infiltration of islets by T cell avatars and by functional assays for insulin secretion. After 18 hours of co-culture, slices showed strong infiltration of ENTPD3^+^ islets by enhanced green fluorescent protein (eGFP) reporter-expressing, IGRP_265-273_ specific T cell avatars and strong labeling of ENTPD3^+^ islets with PPI ImmTAAI (Fig. 7B). Perifusion of slices after 48 hours of co-culture with IGRP-specific avatars showed reductions in glucose- and KCl-induced insulin secretion with ImmTAAI partially rescuing functional beta cell secretion (Fig. 7C); however, the differences were not tested for statistical significance due to a low sample size because of the need to pool multiple slices per perifusion chamber to reach the detection range of the insulin ELISA kit. To improve sample size and throughput, we shifted to testing insulin secretion in single slices using static glucose and KCl incubation. We also shifted to using the PPI_6-14_-specific T cell avatars. As before, we observed eGFP reporter-expressing T cell avatars interacting with islets in slices, with strong labeling of islets with PPI ImmTAAI after the first 18 hours of co-culture (Fig. 7D). After 48 hours of co-culture, the T cell avatars caused a significant reduction in baseline insulin secretion in 3 mM glucose and significant loss of insulin secretion following 16 mM high glucose and KCl stimulation, while ImmTAAI-treated slices retained their function (Fig. 7 E,F). These observations were consistent across pooled data from repeat experiments in three non-diabetic HLA-A2-positive slice donors (nPOD 6637, 6639, 6640; Table S1) (Fig. 7F).

**Fig. 7.**
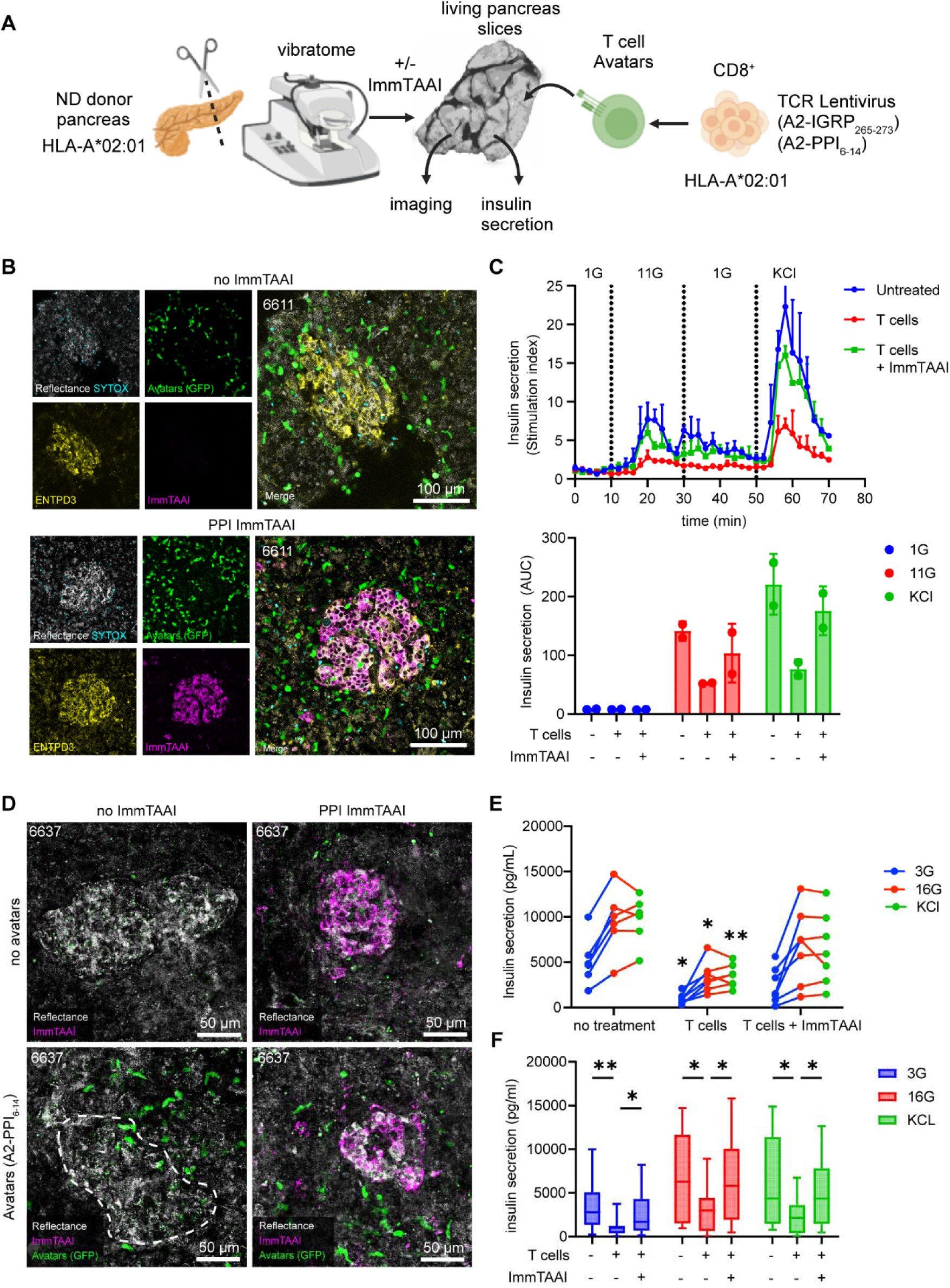
PPI ImmTAAI suppressed killing of beta cells in slices by T cell avatars. (**A**) Experimental scheme depicts the co-culture assay system used to evaluate the immunogenicity of pancreas slices treated with PPI ImmTAAI in the presence of IGRP_265-273_-reactive (Clone 32), or PP_I6-14_-reactive (clone 15b) CD8^+^ avatars. (**B**) Live-cell confocal microscopy images 18 hours after addition of 200,000 IGRP-specific T cell avatars per slice. (**C**) Dynamic perifusion-based glucose and KCl-stimulated insulin secretion and area-under-the-curve (AUC) quantification of slices 48 hours after addition to IGRP-specific T cell avatars with and without PPI ImmTAAI treatment (20 nM). n=2 perifusion chambers with three slices per chamber; mean +/- sem. (**D**) Live-cell confocal microscopy images 18 hours after addition of 200,000 PPI-specific T cell avatars per slice. (**E**) Static glucose and KCl-stimulated insulin secretion from slices 48 hours after addition to PPI-specific T cell avatars with and without PPI ImmTAAI treatment (20 nM). n=6-7 slices per donor. Data are representative of experiments conducted in 3 donors. (**F**) Combined static glucose and KCl-stimulated insulin secretion from 3 donors. Mean +/- range. Statistical differences were determined by two-way ANOVA followed by Tukey’s post-hoc analysis. *p<0.05, ** p < 0.01.

These results establish co-culture of pancreas tissue slices with T cell avatars as an approach to model T1D in tissues from donors without diabetes that creates a condition of autoreactive T cell infiltration of the pancreatic islets and exocrine tissue alongside functional impairment of beta cells. We used this model as an approach to test ImmTAAI as an immunoprotective therapy and showed that it preserves insulin secretory function.

## DISCUSSION

This study investigated the potential for PD-1 agonist ImmTAAI molecules to bind target pHLA in human pancreata and protect against islet autoreactive T cell activity. It was previously shown that cell surface binding of PD-1 agonist ImmTAAI is essential for its activity, as only target cell-bound ImmTAAI molecules can inhibit T cell cytokine secretion and target cell killing (*29*). However, since both the PD-1 agonist nanobody and TCR targeting domain of the ImmTAAI are human-specific, the reagents are not compatible with studies in mice. Here, we used pancreas slice technology to characterize ImmTAAI binding in primary human pancreatic tissue.

Live pancreas tissue slices have been used to study islet biology, beta cell dysfunction, and diabetes pathophysiology (*30, 31, 33, 59–61*). Though slices are not a perfect model of the *in vivo* environment, they offer many advantages over isolated islets (*62*). Our results provide valuable insight into the potential for beta cell-specific ImmTAAI molecules to bind to the intended target cell in heterogeneous, complex tissue environments such as those found *in vivo*. We showed that PPI ImmTAAI molecules bind specifically to beta cells in a dose-dependent manner. Importantly, labeling ImmTAAI with a CF647 fluorophore did not affect its binding kinetics or potency in cell-based assays, providing confidence that the binding we observed in tissue slices accurately reflects what would occur with unlabeled ImmTAAI.

A key piece of data presented here demonstrates ImmTAAI binding in a pancreas actively undergoing autoimmune attack, as nPOD donor 6578 had a substantial number of islets with active insulitis defined as at least three islets per pancreas section with ≥15 CD45^+^ cells or ≥6 CD3^+^ T cells immediately adjacent to or within the islet (*63, 64*), at the time of organ collection, as documented on the nPOD histopathology database (https://www.jdrfnpod.org/for-investigators/online-pathology-information/). It is likely that pancreata with active autoimmunity and remaining beta cell mass can bind significant amounts of ImmTAAI since HLA class I hyper-expression on beta cells is a characteristic feature of T1D (*65–67*). Additionally, inflammation causes increased beta cell PD-L1 expression (*68, 69*), and we showed previously that PD-1 agonist ImmTAAI molecules are additive with cellular PD-L1 in triggering PD-1 activity on T cells (*29*). This evidence, taken together with the data shown here, suggests that targeting HLA-A2 presented PPI_15-24_ is a viable strategy for inducing immune tolerance towards beta cells by decreasing CD8^+^ T cell-mediated destruction through activation of the PD-1 pathway in islet-infiltrating T cells.

Although it is critical to perform studies in T1D and AAb+ pancreas donors, these donors are limited and heterogeneous. Thus, we devised an approach to carry out studies in human tissue in which we can control the timing and degree of autoimmunity. To achieve this, we simulated the autoimmune environment of T1D by introducing engineered T cell avatars into living pancreas slices from histocompatible non-diabetic donors. The T cell avatars expressed TCRs specific to beta cell antigens IGRP and PPI. We found that the presence of the islet antigen reactive T cells was sufficient to cause beta cell dysfunction and/or loss. ImmTAAI treatment preserved beta cell function in this model system, further supporting its future clinical evaluation.

### Limitations of the study

The sparse nature of pancreatic organ donor material limits our capacity to obtain living pancreatic slices, including the need for HLA matching to ImmTAAI molecules and T cell avatars. The functional measurements conducted in slice co-cultures with T cell avatars were not directly quantified for the number of surviving beta cells. Although slices contain intact vasculature, this vasculature is not perfused during our experiments. Thus, the cues and mechanisms by which T cells enter slice tissues may differ from that occurring in vivo. Although HLA-A2 is one of the most frequent MHC class I alleles among T1D patients (*70*), the specific PPI_15-25_ ImmTAAI molecule described in this study would require modification with a different TCR targeting domain to be effective for other HLA alleles.

### Translation to the clinic

Our approach has the advantage of being a soluble protein that is engineered to be highly specific for beta cells through the paired TCR structure and has no activity when unbound (*29*). This class of bi-specific, tissue-targeted drugs has recently seen success in cancer therapy, with one bi-specific intended to treat metastatic uveal melanoma completing a Phase III clinical trial and receiving United States Food and Drug Administration (FDA) approval in 2022 (*71*). The next step for translation of this class of tissue-targeted PD-1 agonist is a safety, dosing, and efficacy trial to preserve endogenous beta cell mass in T1D without the need for systemic immune suppression.

## MATERIALS AND METHODS

### Experimental Design

#### Research objectives

The objectives of this study were to determine if ImmTAAI molecules specifically bind to beta cell surfaces in native tissue environments and whether ImmTAAI molecules inhibit cellular immunity in type 1 diabetes.

#### Research subjects or units of investigation

Organotypic slices from human pancreas and cultured cells.

#### Experimental design

Controlled laboratory experiments with PPI ImmTAAI compared to off-target Tel ImmTAAI and untreated samples. Measurements of ImmTAAI molecule localization were by confocal microscopy and T cell effector function and target cell killing were by *in vitro* biochemical assays.

#### Randomization

Individual pancreas slices were assigned to treatment conditions randomly.

#### Blinding

Researchers were blinded to the treatment condition during confocal image acquisition and image quantification. *In vitro* biochemical assays were performed unblinded but in multi-well plate format where all samples were treated the same.

#### Sample size

For human pancreas slices, sample size was determined by the availability of HLA-A2^+^ donor organs, which are intrinsically limited in number, particularly for at-onset T1D donors. Studies in human tissues were validated by parallel *in vitro* assays with cells / cell lines.

#### Rules for stopping data collection

Data was collected for the total available tissue donors that met our selection criteria over the years 2021-2025 in accordance with the contract agreement between Immunocore and the University of Florida.

#### Data inclusion/exclusion criteria

Currently, nPOD sets organ selection criteria for recovery of pancreata from T1D donors who are younger than 30 years of age and with T1D duration ≤ 7 years. These selection criteria are based on years of data that residual insulin-positive islets and insulitis are more likely to be found using these acceptance criteria. Control donors are defined as BMI<30, aged 0-40 years with no diagnosis of diabetes. Donors were excluded from the study if the slices had low viability (greater than 10% dead cells) or poor islet function (insulin secretion stimulation index less than 2, except for T1D donors, which are expected to have poor islet function). All donors in the study expressed HLA-A2, except for the negative controls for HLA type.

#### Outliers

No outliers were excluded from the data.

#### Selection of endpoints

Not applicable to this study.

#### Replicates

In vitro ImmTAAI activity experiments with cell lines (Figs. 1,4) were repeated three times. Experiments to optimize the binding conditions (concentration, incubation time) for ImmTAAI molecules in human pancreas slices were conducted using multiple slices from one or two donors per optimization parameter (ImmTAAI TCR specificity, concentration, incubation time, and donor HLA-type) (Figs. 2,3,S1), while ImmTAAI molecule binding at the optimal conditions were repeated in 16 donors (Table S1). ImmTAAI activity in T1D pancreas was conducted in a single, rare HLA-A2^+^ organ donor with at-onset T1D and active insulitis (nPOD 6578) (Fig. 5). *In vitro* ImmTAAI cell killing experiments with T cell avatars (Fig. 6) were repeated twice using two different sBC stem cell lines. ImmTAAI activity to preserve islet function in non-diabetic tissue with the addition of T cell avatars was performed in a single HLA-A2^+^ pancreas for dynamic insulin secretion (6611) and three donors for static insulin secretion (6637, 6639, 6640) (Fig. 7).

### ImmTAAI production

PPI ImmTAAI was expressed as a soluble molecule in ExpiCHO-S cells (Thermo Fisher Scientific) and purified using three stages of chromatography: anion, cation, and size exclusion (*29*). A control molecule, which binds pHLA from the tumor-associated human telomerase reverse transcriptase (hTERT_540–548_) that is not presented by human cancerous or normal cells (*72*) (Tel ImmTAAI), was expressed as inclusion bodies in E. coli followed by refolding and purification using anion and size exclusion chromatography (*73*). CF647 succinimidyl ester labelling was performed according to the manufacturer’s recommendations (Biotium).

### SPR kinetic analysis

SPR binding analysis was performed using a BIAcore 8K (GE Healthcare) as described previously (*29*). Briefly, biotinylated pHLA-A2 complexes were immobilized on a streptavidin-coated CM5 chip. ImmTAAI was injected over the immobilized pHLA using single cycle kinetics to assess pHLA affinity. Assessment of PD-1 binding was performed using pHLA-immobilized ImmTAAI followed by injection with PD-1. Data were analyzed using BIAanalysis software (Biacore Insight Evaluation software version 3.0.12.15655, GE Healthcare).

### Cellular TCR signaling assay using the Jurkat NFAT luciferase PD-1 reporter

The ECN90 Jurkat NFL Mel5 PD-1 reporter assay was performed as described previously (*29*). ECN90 cells were plated at 50,000 cells/well in Optiβ3 media into a white 96-well culture plate pre-coated with β-coat and incubated for 16-20 hours at 37°C, 5% CO_2_. Melan-A_26-35_ peptide (ELAGIGILTV heterolytic peptide) was added for 2 hours followed by titrations of ImmTAAI molecules. The assay was initiated by adding 50,000 Jurkat NFL Mel5 PD-1 effector cells and incubating for 16 – 20 hours. For assessment of selectivity, ECN90 cells were replaced with NCI-H1703 cells pulsed with 5 µM Melan-A peptide and treated in an identical fashion. The reporter assays were developed by adding BioGlo reagent (Promega) with NFAT activity determined by measuring luciferase luminescence on a plate reader (Clariostar, BMG Labtech). NFAT activity was normalized against TCR-stimulated controls, with ImmTAAI dose response data analyzed in GraphPad Prism using a four parameter, non-linear least squares fit to determine IC_50_ values.

### Pancreatic organ donors

Pancreas tissue slices were obtained through the Network for Pancreatic Organ donors with Diabetes (nPOD, RRID: SCR_014641, Gainesville, Florida; https://www.jdrfnpod.org) from donors without diabetes (n=10), donors who were single GAD autoantibody positive (AAb^+^) without diabetes (n=3), and donors with recent-onset T1D (n=2, All procedures using human slices were performed according to the established standard operating procedures (SOP) of the nPOD Organ Processing and Pathology Core (OPPC) as approved by the University of Florida Institutional Review Board (IRB201600029) and the United Network for Organ Sharing (UNOS) according to federal guidelines, with informed consent obtained from each donor’s legal representative. For each donor, a medical chart review was performed and C-peptide measured, with T1D diagnosed according to the guidelines established by the American Diabetes Association (ADA). Demographic data, hospitalization duration, and organ transport time were obtained from hospital records. Donor pancreata were recovered, placed in transport media on ice, and shipped via organ courier to the University of Florida. The tissue was processed by a licensed pathology assistant. nPOD tissues used for this project were approved as nonhuman by the University of Florida IRB (NH00041892, NH00042022).

S1). All procedures using human slices were performed according to the established standard operating procedures (SOP) of the nPOD Organ Processing and Pathology Core (OPPC) as approved by the University of Florida Institutional Review Board (IRB201600029) and the United Network for Organ Sharing (UNOS) according to federal guidelines, with informed consent obtained from each donor’s legal representative. For each donor, a medical chart review was performed and C-peptide measured, with T1D diagnosed according to the guidelines established by the American Diabetes Association (ADA). Demographic data, hospitalization duration, and organ transport time were obtained from hospital records. Donor pancreata were recovered, placed in transport media on ice, and shipped via organ courier to the University of Florida. The tissue was processed by a licensed pathology assistant. nPOD tissues used for this project were approved as nonhuman by the University of Florida IRB (NH00041892, NH00042022).

### Pancreas slice generation and culture

Pancreas tissue slices were generated by nPOD in accordance with published methods (*33*). Pieces of approximately 1 g were obtained from the pancreas body-tail juncture and placed into extracellular solution (ECS) (125 mM NaCl, 2.5 mM KCl, 26 mM NaHCO_3_, 1.25 mM NaH_2_PO_4_, 1 mM MgCl_2_, 2 mM CaCl_2_, 10 mM HEPES, 3 mM glucose, pH 7.4) supplemented with 25 kIU/ml aprotinin (Sigma #A6106). These were cut into smaller tissue blocks, and connective, adipose, and fibrotic tissues were removed. Small tissue blocks of about 0.5 cm^3^ were embedded in low-melting-point agarose (3.8%) and mounted on a specimen holder using superglue (90-120 cps, World Precision Instruments). The holder was connected to the tray of a semiautomatic vibratome (Leica VT1200S) and filled with cold ECS supplemented with aprotinin. Tissue slicing was performed at a step size of 120 μm, speed of 0.1 mm/s, amplitude of 1 mm, and an angle of 15 degrees. Slices were collected into a 100-mm Petri dish containing 3 mM glucose buffer (137 mM NaCl, 5.36 mM KCl, 0.34 mM Na_2_HPO_4_, 0.81 mM MgSO_4_, 4.17 mM NaHCO_3_, 1.26 mM CaCl_2_, 0.44 mM KH_2_PO_4_, 10 mM HEPES, 0.1% BSA, 3 mM glucose, pH 7.3) supplemented with aprotinin. Slices were cultured in low-glucose DMEM containing 10% FBS, 1:100 antibiotic antimycotic solution (Corning), and aprotinin in an incubator at 24°C, 5% CO_2_.

### Slice Fluorescent staining and imaging

Slices were washed twice with pre-warmed (37°C) Kreb’s ringer bicarbonate HEPES buffer (KRBH, 115 mM NaCl, 4.7 mM KCl, 2.5 mM CaCl_2_, 1.2 mM KH_2_PO_4_, 1.2 mM MgSO_4_, 25 mM NaHCO_3_, 25 mM HEPES, 0.2% BSA, 3 mM glucose) and transferred to a μ-Slide 8 well, chambered coverslip (Ibidi) with one slice per well containing 250 μl pre-warmed KRBH with ImmTAAI (0, 2, or 20 nM) and 0.5 μl Alexa Fluor 594-labeled anti-ENTPD3 at 37°C, 5% CO_2_ for one or two hours, as indicated, washed with KRBH buffer 3 times, then labeled with 1 μM SYTOX Blue viability stain (Thermo Fisher Scientific) and positioned in the chambered coverslips under a slice anchor (Warner Instruments) for imaging on a Leica SP8 or Leica Stellaris confocal microscope.

### Killing and motility assay

EndoC-βH2 mKate red were generated by transducing EndoC-βH2 cells (Human Cell Design) with HLA-A201/β2-microglobulin and mKate red lentivirus (NucLight red, Sartorius) as described (*29*). EndoC-βH2 cells were plated at 5×10^4^ cells per well on TPP-96-well plate pre-treated with β-coat solution (Human Cell Design) in Ultiβ1 media and incubated over night at 37°C, 5% CO_2_. The cells were pulsed with PPI_6-14_ peptide at 2 μM for 2 hours. ImmTAAI molecules were added at 10 nM and incubated for 2 hours. PPI_6–14_-HLA-A2-specific autoreactive T cell clone 4b PD-1+ was labelled with Cell Trace Violet (eBioscience) and added to the plate. Co-culture plates were live imaged by Opera Phenix Plus confocal microscope (PerkinElmer) at 37 °C and 5% CO_2_. All conditions were performed in triplicate. For the motility assay, the β-cells and T cells were plated at 1:4 effector:target ratio and the images acquired with 40X water objective, 8 fields/well with iterations every 2.36 minutes for 2 hours. For the killing assay, the cells were plated at 1:1 effector:target ratio and images acquired using 10X air objective, 4 fields/well, with iterations every 2 hours for 72 hours. Images were analyzed using Harmony Software (Perkin Elmer).

### Generation of T cell avatars

Fresh peripheral blood mononuclear cells (PBMCs) were obtained from human leukapheresis-enriched blood of healthy donors (median age: 31.5 years, range 19-38 years, *N*=4, 75% male) purchased from LifeSouth Community Blood Centers (Gainesville, FL, USA). Lentiviral transduction was used to generate HLA-A*02:01 restricted CD8^+^ T cell avatars that recognize islet-specific glucose-6-phosphatase catalytic subunit (IGRP_265-273_) (*51*), melanoma antigen (Melan-A; MART-1) (*54*), pre-proinsulin (PPI_6-14_; clone 15b), or express an HLA-A2 directed chimeric antigen receptor (HLA-A2-CAR) (*55*), as previously described (*50*). Briefly, peripheral blood was overlayed onto Ficoll-Paque Plus medium (Thermo Fisher) for density gradient centrifugation (1200 x g, 20 minutes). Recovered PBMCs were treated with Ammonium-Chloride-Potassium (ACK) Lysis Buffer (Gibco) for 5 minutes at 4°C before quenching with 1x PBS. Naïve CD8^+^ T cells were isolated from the PBMC fraction using the EasySep™ Human Naïve CD8^+^ T Cell Isolation Kit (STEMCELL Technologies) according to the manufacturer’s instructions.

Naïve CD8^+^ T cells were plated at 2.5 x 10^5^ cells/well in 1 mL of complete RPMI media (cRPMI; RPMI 1640 media Phenol Red w/o L-Glutamine, 5 mM HEPES, 5 mM MEM Non-Essential Amino Acids (NEAA), 2 mM Glutamax, 50 µg/mL penicillin, 50 µg/mL streptomycin, 20 mM sodium pyruvate, 50 mM 2-mercaptoethanol, 20 mM sodium hydroxide, and 10% FBS, supplemented with 100 IU/mL rhIL-2 and 5 ng/mL rhIL-7. The cells were stimulated with Dynabeads™ Human T-Expander CD3/CD28 (Thermo Fisher) at a 1:1 bead:cell ratio for 48 hours, after which the cells were treated with protamine sulfate (8 µg/ml) and 3 transducing units (TU)/cell of either the IGRP, MART-1, HLA-A2-CAR, or PPI lentiviral vector before spinnoculation (1000 g x 30 minutes at 32°C). Cell culture media was changed every 3-4 days, and beads were removed on day nine.

After expansion, cells were cultured in cRPMI supplemented with 100 IU/mL rhIL-2 for three days before enriching for a pure population of successfully transduced, GFP^+^ T cell avatars, using fluorescence-activated cell sorting (FACS), accomplished with a FACSMelody™ Cell Sorter (Beckton Dickinson).

### Chromium-release assay

^51^Cr-Release assays were performed as described (*56*) to assess the cytotoxic capacity of T cell avatars in the presence of ImmTAAI. Briefly, 30,000 sBC were seeded in Cultrex-coated wells in a 96-well plate and pre-treated for two days with IFN-γ (100 ng/ml) in CMRL media containing 5% N-21 MAX, 1% NEAA, 1% GlutaMAX, 10 ug/ml Heparin, 2mM Cysteine, 10 μM Zinc, 100 μM BME, 1 μM T3, 10 μM Alk5i II RepSox, 50 ug/ml VitC, 0.1% Trace Elements A, 0.1% Trace Elements B, and 1x Penicillin-Streptomycin. sBCs were loaded with radiolabeled ^51^CrNa_2_O_4_ (Revvity) at an activity of 1.48 x 10^5^ Bq/well for four hours in DMEM, 5 mM HEPES, 5 mM MEM NEAA, 50 µg/ml penicillin, 50 µg/ml streptomycin, 0.02% Bovine Serum Albumin, and 10% FBS, and washed twice with fresh media. Following radiolabeling, sBCs were co-cultured with MART-1, PPI, IGRP, or HLA-A2-CAR T cell avatars at 0:1, 1:1, 5:1, and 10:1 effector:target ratios for 48 hours. For co-cultures performed in the presence of ImmTAAI, unlabeled ImmTAAI was added at a concentration of 20 nM at the start of the co-culture. Following 24 hours of co-culture for IGRP avatars or 48 hours of co-culture for PPI avatars, the supernatants were removed and transferred into 6×50 mm lime glass tubes. The lysates of adherent cells were collected using a 2% SDS wash and transferred into separate tubes. ^51^Cr activity, measured in counts per minute (CPM), was assessed for both fractions on a Wizard 1470 automatic gamma counter (Revvity). The specific lysis of sBCs was calculated as follows:

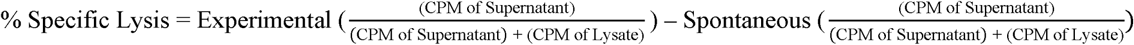

### Cytokine multiplex assay

To quantify the production of cytokines by T cells after co-culture with sBCs in the presence and absence of ImmTAAI, cell culture supernatants from the CML assay were used to evaluate IL-2, IL-4, IL-6, IL-10, IL-17A, FasL, IFN-γ, TNF, granzyme A, granzyme B, perforin, and granulysin production. Using the LEGENDplex™ Human CD8/NK Panel Kit (BioLegend) and a 5L Aurora spectral flow cytometer. Data were analyzed using the LEGENDplex Data Analysis Software Suite (version 2024-12-24; BioLegend). Dilution factors and analyte detection ranges are described in Table S3.

### Co-culture of T cell avatars with pancreas slices

Pancreas tissue slices were maintained in DMEM, 10% FBS, 25 kIU/mL aprotinin from bovine lung (Sigma #A6106), and 1% antibiotic-antimycotic solution (Corning #30-004-CI) at 24°C. Slices were transferred to 12-well plates, 1 slices per well, and equilibrated at 37°C for 1 hour prior in 2 ml modified slice culture media without phenol red, without aprotinin, and supplemented with rhIL-7 and rhIL-15 at 10 ng/ml. ImmTAAI molecule 20 nM was preincubated for 4-5 hours prior to addition of T cell avatars. Subsequently, Corning™ PYREX™ cloning cylinders (Thermo Fisher #0955221) were positioned on top of slices, and 2×10^5^ T cell avatars (IGRP specific or PPI specific) were added as a 50 μl cell suspension (in modified slice media) pipetted slowly into the cloning cylinder. After 2 hours of incubation, cloning cylinders were removed, and the co-culturing system was incubated for 48 hours with live imaging every 24 hours.

### Glucose-stimulated insulin secretion (GSIS) of pancreas slices

Perifusion studies were conducted using a Biorep Technologies Perifusion System (Biorep Technologies) as described (*74*). Three slices each were placed into three closed perifusion chambers and connected to the system. Tissue slices were perifused at a flow rate of 100 μl/min at 37°C with DMEM media supplemented with 3.2 g/l sodium bicarbonate, 1.11 g/l HEPES, 0.11 g/l sodium pyruvate, 4 mM L-glutamine, pH 7.4, 0.1% BSA, 25 kIU/ml aprotinin and 1 mM glucose, 11 mM glucose, or 30 mM KCl. The perifusate was collected in 96-well plates at 2-minute intervals. Perifusates were stored at -80°C until commercial insulin ELISAs were run (Alpco # 80-INSHU-CH01). Static GSIS was performed immediately after the co-culturing experiments. GSIS was performed in 250 μl KRBH supplemented with glucose at 3 mM glucose, 16.7 mM glucose, or 30 mM KCl. Each stimulation was incubated for 1 hour at 37°C sequentially. Each slice was washed once with 500 μl KRBH prior to addition of the next stimulation. The stimulation supernatants were collected, centrifuged at 500 x g for 5 minutes, and stored at -80°C until insulin measurements by ELISA kit (Alpco # 80-INSHU-CH01).

### Statistical Analysis

Sample size (n) was defined as the number of separate islets measured per condition. Means among three or more groups were compared by one-way or two-way analysis of variance (ANOVA) followed by Tukey’s post-hoc pairwise comparisons using GraphPad Prism 9 software. A confidence level of 95% was considered significant. Statistical test used, exact *P* values, error bars, and definition of *n* are all indicated in the individual Fig. legends.

## List of Supplementary Materials

Supplementary Materials and Methods

Fig. S1 to S4

Table S1 to S4

References (*75–82*) (numbers for references only cited in SM)

## Acknowledgments

The authors thank the nPOD donors and their families for the gift of tissues. We thank Amanda Posgai (University of Florida Diabetes Institute) for editing the manuscript, Drew Bloss (University of Florida Diabetes Institute) for technical assistance with slice perifusion, and Helmut Hiller, Maria Beery, Ellen Verney, and Irina Kusmartseva (nPOD Organ Processing and Pathology Core Staff, University of Florida) for slice generation.

## Author contributions

Conceptualization: PW, HA, TMM, TMB, GB, EAP Methodology: MWB, MB, MKH, JMB, TMM, TMB, GB, EAP

Investigation: MWB, MB, KW, ALC, MKH, AEC, PS, SMF, DS, JMB, AML, DMD, MAA, EAP

Visualization: MWB, MB, GB, EAP

Funding acquisition: MWB, MKH, MAA, HAR, TMB, EAP Project administration: PW, HA, TMB, GB, EAP

Supervision: MAA, PW, HA, HAR, TMM, TMB, GB, EAP

Writing – original draft: MWB, EAP

Writing – review & editing: MAA, HA, TMM, TMB, GB, EAP

## Data and materials availability

All data are available in the main text or the supplementary materials. Availability of ImmTAAI molecules and PPI_6-14_ TCRs for non-commercial, research purposes is subject to approval by Immunocore and requires a materials transfer agreement (MTA).

