## Supplementary Information for "Inhibition of cell-mediated immunity in type 1 diabetes by beta cell-targeted PD-1 agonists in pancreas tissue slices"

### SUPPLEMENTARY FIGURES

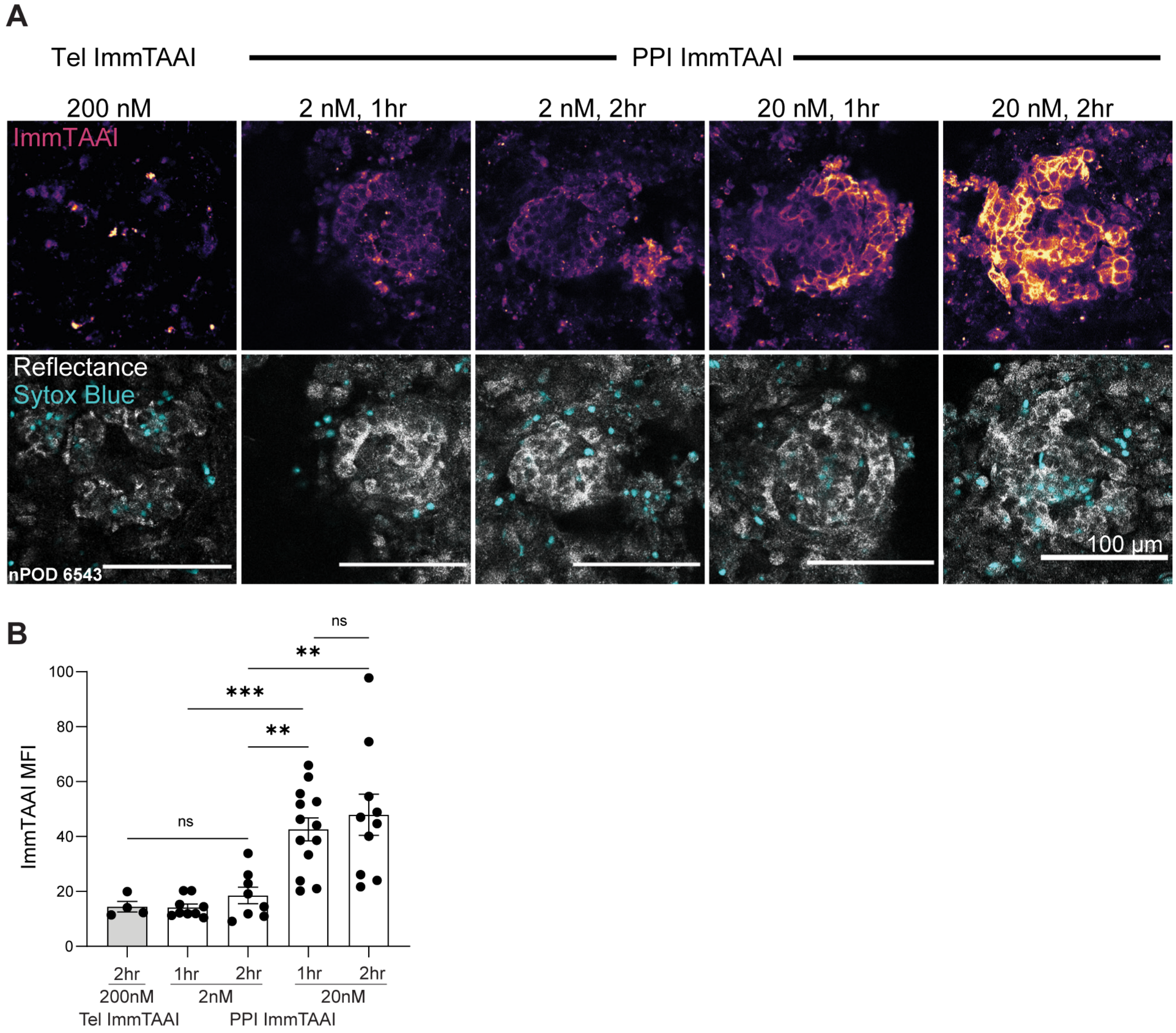


**Supplementary Figure 1. Optimizing ImmTAAI binding conditions to live pancreatic tissue slices.** (**A**) Confocal microscopy images of pancreas slices incubated with CF647-labeled PPI ImmTAAI at 2 or 20 nM for 1 or 2 hours. Slices were also stained with Sytox Blue for viability. (**B**) MFI quantification of microscopy images showing increased PPI ImmTAAI binding at 20 nM, with the highest average MFI at 2 hours. Each dot represents a separate islet from 1 separate slice per condition and is the mean of 2-4 z-planes taken per islet. Statistical differences were determined by one-way ANOVA followed by Tukey's post-hoc analysis. ns = non-significant, ** $p < 0.01$, *** $p < 0.001$.

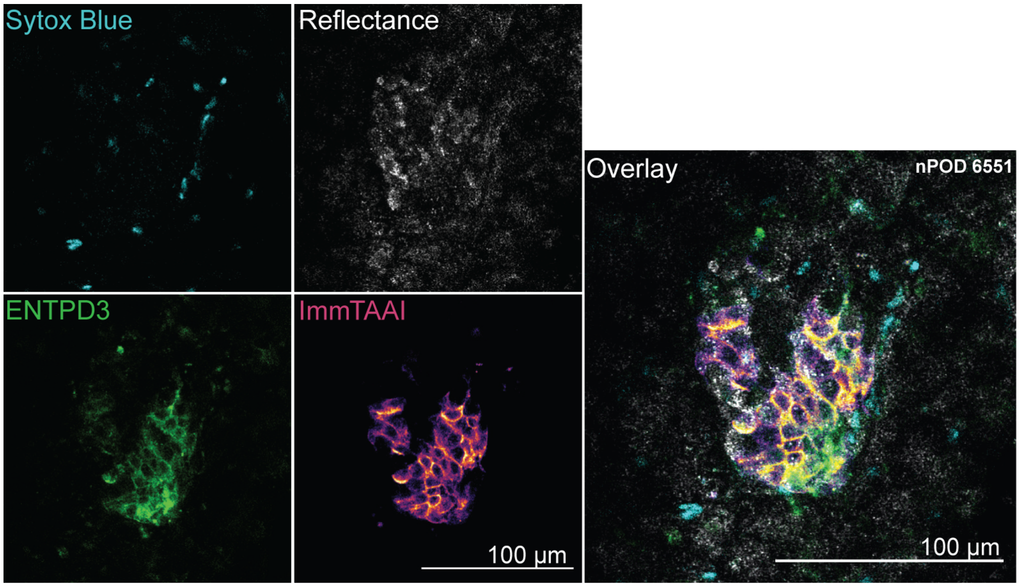


**Supplementary Figure 2. PPI ImmTAAI binding in a donor with recent diagnosis of T1D and residual beta cells.** Live cell confocal microscopy of ImmTAAI binding to ENTPD3$^{+}$ beta cell surfaces in remaining islets in a donor with T1D.

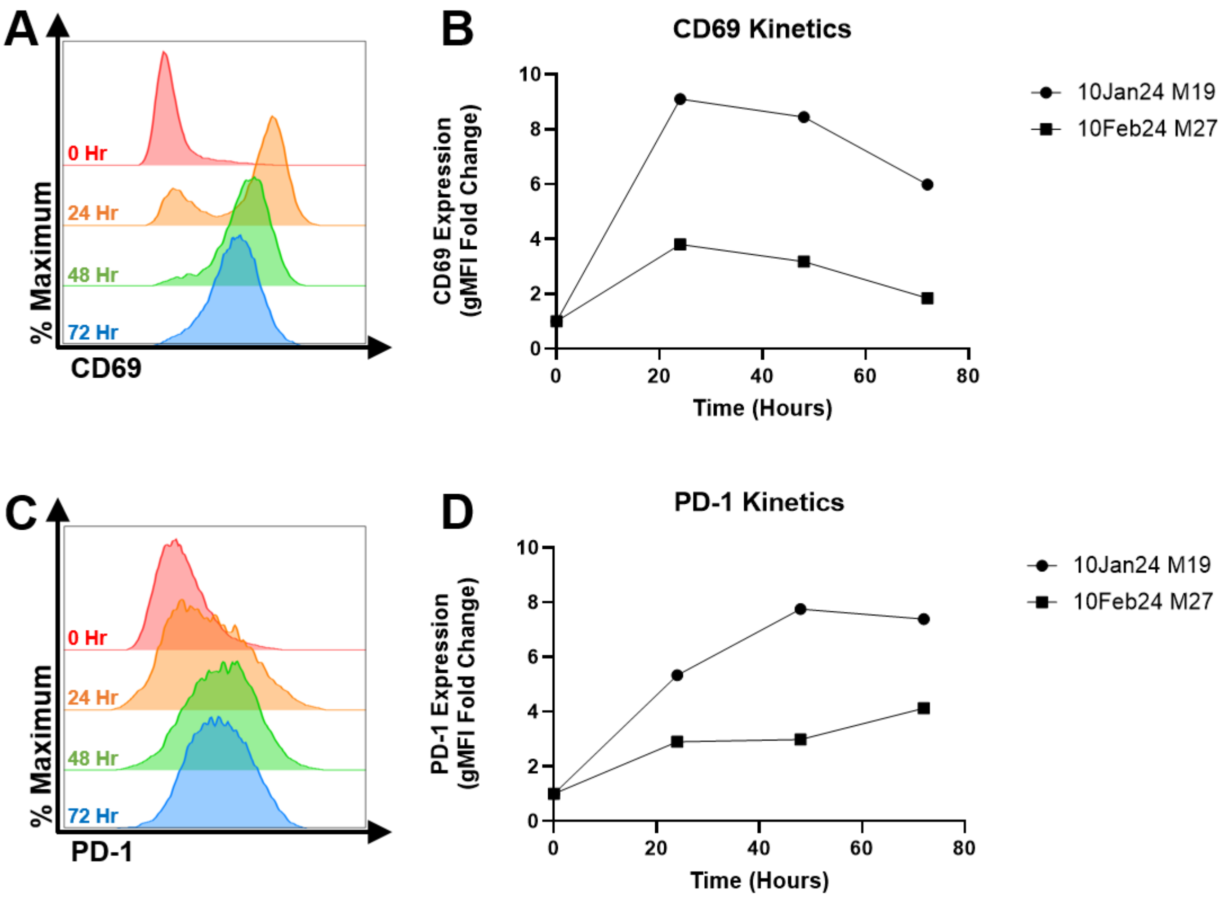


**Supplementary Figure 3. Assessment of avatar activation capacity.** Flow cytometric assessment of (**A-B**) CD69 and (**C-D**) PD-1 expression on expanded IGRP-reactive CD8+ T cell avatars after anti-CD3/CD28 stimulation was used to evaluate functionality. (A, C) Representative histograms show expression of activation markers following 0 (red), 24 (orange), 48 (green), or 72 (blue) hours of stimulation with (B, D) paired dot plots showing the activation marker expression kinetics, relative to baseline. N = 2 biological.

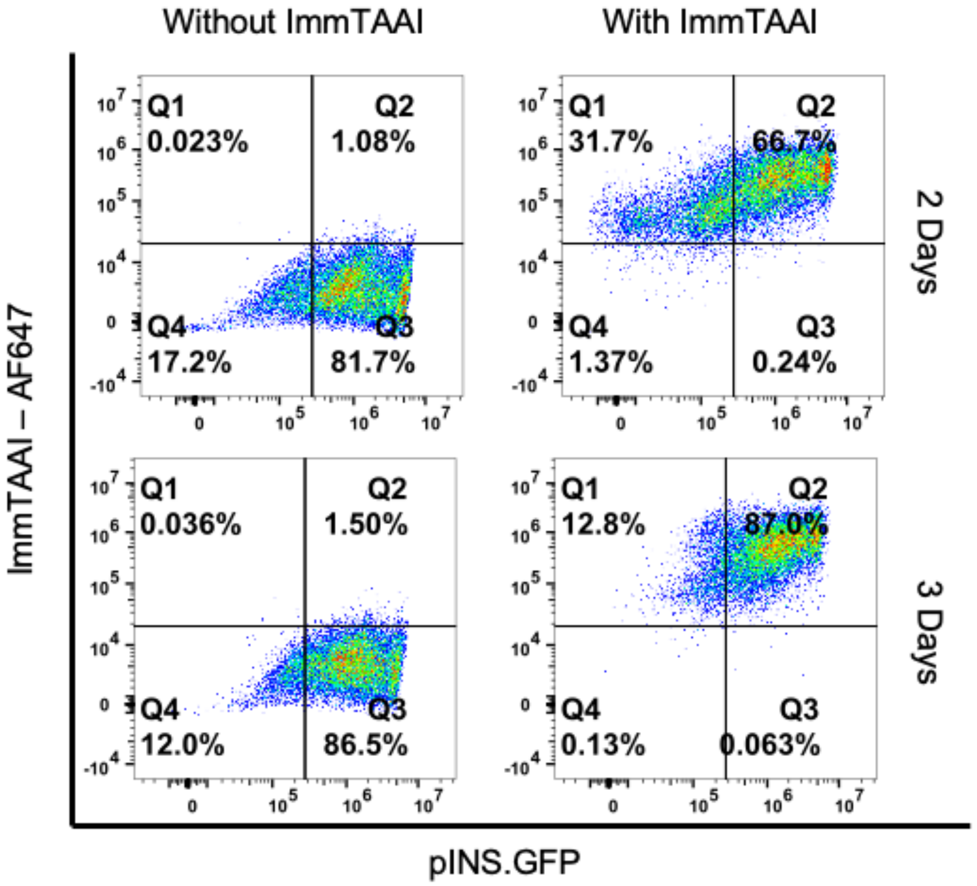


**Supplementary Figure 4. PPI ImmTAAI binding to stem cell-derived beta cells.** Flow cytometric assessment of PPI ImmTAAI binding to stem cell-derived beta cells during reaggregation into clusters on day 2 and day 3 of aggregation. Beta cells express eGFP under the insulin promoter. PPI ImmTAAI added at 20 nM in reaggregation culture media.

**Supplemental Tables:**

**Supplemental Table 1: Donor information.**

| nPOD ID | RRID # | Donor Type | AutoAbs | Age (yrs) | Sex | C-pep (ng/mL) | HLA-A | Experiments used |
|---|---|---|---|---|---|---|---|---|
| 6538 | SAMN25652249 | Aab+ | GADA+ | 19 | Male | 11.33 | 01:01/02:01 | Fig 2A |
| 6539 | SAMN25652250 | No diabetes | Negative | 24 | Male | 39.23 | 02:01/11:01 | Fig 2C-E |
| 6540 | SAMN25652251 | No diabetes | Negative | 6 | Female | 2.03 | 29:01/31:01 | Fig 3A-C |
| 6543 | SAMN25652254 | No diabetes | Negative | 3 | Male | 1.45 | 02:01/31:01 | Fig S1 |
| 6548 | SAMN25652259 | No diabetes | Negative | 20 | Male | 4.04 | 02:01/30:02 | Fig 2C |
| 6551 | SAMN25652262 | T1D (0.5 yrs duration) | GADA+ IA2A+ mIAA+ ZnT8A+ | 20 | Male | 0.11 | 02:01/29:02 | Fig S2 |
| 6552 | SAMN30386842 | No diabetes | Negative | 33 | Female | 1.8 | 31:01/32:01 | Fig 3A-C |
| 6553 | SAMN30386843 | Aab+ | mIAA+ | 12 | Female | 4.62 | 02:01/03:01 | Fig 3A-C |
| 6575 | SAMN33284293 | Aab+ | GADA+ | 23 | Male | 5.21 | 01:01/02:01 | Fig 3F-G |
| 6578 | SAMN33284295 | T1D (0.0 yrs duration) | IA2A+ ZnT8A+ | 11 | Female | 0.35 | 02:01/03:01 | Fig 4A-D |
| 6611 | SAMN44486497 | No diabetes | Negative | 14 | Male | 10.98 | 02:01/02:01 | Fig 2B, Fig 6B |
| 6615 | SAMN44486501 | No diabetes | Negative | 14 | Male | 2.9 | 02:01/68:01 | Fig 3D |
| 6637 | SAMN49972145 | Aab+ | GADA+ | 36 | Male | 14.36 | 02/29 | Fig 6D,E |
| 6639 | Pending | No diabetes | Negative | 12 | Female | 9.13 | 02/02 | Fig 6E |
| 6640 | Pending | No diabetes | Unavailable (no serum) | 4 | Female | Unavailable | 02/24 | Fig 6E |

**Supplemental Table 2: Antibodies Used for Flow Cytometry**

| Target | Clone | Fluorophore | Vendor | Host Species | Concentration | RRID |
|---|---|---|---|---|---|---|
| CD8 | SK1 | Alexa Fluor 700 | BioLegend | Mouse | 1.00 µg/mL | AB_2562790 |
| CD69 | FN50 | BV421 | BioLegend | Mouse | 1.00 µg/mL | AB_2561909 |
| PD-1 | EH12.2H7 | BV650 | BioLegend | Mouse | 1.00 µg/mL | AB_2566362 |

**Supplemental Table 3: LEGENDplex Detection Ranges**

| LEGENDplex Assay Supernatant Dilutions | | |
|---|---|---|
| Analyte | Dilution Factor | Adjusted Detection Range (pg/mL) |
| IL-17A | 1; ND | 2.9 to 12,000 |
| IL-2 | 1; ND | 16.6 to 68,000 |
| IL-4 | 1; ND | 3.9 to 16,000 |
| IL-10 | 1; ND | 3.7 to 15,000 |
| IL-6 | 1; ND | 4.6 to 19,000 |
| TNF | 1 | 3.9 to 16,000 |
| Fas | 1; ND | 14.9 to 61,000 |
| FasL | 1 | 2.4 to 10,000 |
| IFN-γ | 100 | 39 to 160,0,00 |
| Granzyme A | 2 | 12.6 to 52,000 |
| Granzyme B | 2 | 28.2 to 116,000 |
| Perforin | 1 | 2.9 to 12,000 |
| Granulysin | 1 | 11.0 to 45,000 |
| ND = Not Detectable; Lot: B435969 | | |

### SUPPLEMENTARY MATERIALS AND METHODS

**High-resolution HLA typing.** HLA typing was carried out according to nPOD protocols (*75*). Genomic DNA was extracted from frozen spleen tissue samples of every donor using DNeasy Blood & Tissue kit (Qiagen). The quality and concentration of purified DNA were assessed by spectrometry using an Epoch microplate absorbance reader (Agilent). High-resolution HLA typing was performed at the Autoantibody/HLA Core Facility, Barbara Davis Center for Childhood Diabetes (Aurora, Colorado). Briefly, the specific HLA region to be typed was amplified, labeling the amplicon with Biotin. Amplicon was applied to sequence-specific oligonucleotide-coupled microspheres (One Lambda). After incubation, Streptavidin-PE was added to the bead/amplicon mixture. Microspheres were visualized using a LabScan3D instrument (Luminex). The HLA alleles were determined by HLA Fusion software (One Lambda, Los Angeles, California).

**Cell lines.** The Jurkat NFAT luciferase T cell line (Promega) was maintained in RPMI-1640, 10% FBS, 2 mM L-glutamine, 0.1 mM MEM non-essential amino acids, 1 mM sodium pyruvate, 50 U/ml penicillin, 50 μg/ml streptomycin and 200 μg/ml hygromycin B. The immortalized human pancreatic beta cell lines, ECN90 and EndoC-βH2 (Human Cell Design) were maintained in Optiβ3 and Ultiβ1 media, respectively, in tissue culture vessels pre-coated with β-coat (Human Cell Design). The NCI-H1703 non-small lung carcinoma cell line (ATCC) was maintained in RPMI-1640, 10% FBS, 2 mM L-glutamine, 50 U/ml penicillin, and 50 μg/ml streptomycin. Cells were maintained at 37°C, 5% $CO_2$.

**Lentivirus production and transduction.** Lentiviruses containing PDCD1 (PD-1, obtained from Origene, Rockville, Maryland) or the Mel5 TCR specific for Melan $A_{26\text{-}35}$ peptide (produced in-house) presented by HLA-A*02 were packaged by transfection of lentiviral vectors with packaging plasmids (pMD2.g, pMDLg/p RRE and pRSV.REV, in-house) into HEK293T cells with TurboFect™ Transfection Reagent (Thermo Fisher Scientific, Waltham, Massachusetts). Lentiviral particles were collected 48 hours after transfection, filtered through a 0.45 μm filter and concentrated by centrifugation at 10,000 x g for 16 hours. Pellets were re-suspended in growth media and stored at -80°C prior to use. Lentivirus was added to 1 x $10^6$ exponentially growing Jurkat NFAT luciferase T cells in wells of a 24-well cell culture plate. After 48 to 72 hours, transduced cells were harvested, assessed for expression of transduced genes by flow cytometry and expanded into cell culture flasks.

**ENTPD3 antibody labeling.** Human anti-ENTPD3 antibody (#AF4400, R&D Systems, Minneapolis, Minnesota) was reconstituted to 2 mg/ml in sterile 0.1 M $NaHCO_3$ buffer and labeled with AlexaFluor 594 NHS ester (Invitrogen, Waltham, Massachusetts) according to the manufacturer's recommendations. Unbound dye was removed by a Zeba 0.5 mL desalting column with 7 kDa molecular weight cut-off (Thermo Fisher Scientific, Waltham, Massachusetts). Purified, labeled antibody was aliquoted and stored at -20°C until use.

**Image analysis.** Mean fluorescent intensity (MFI) and co-localization were quantified using Fiji software (ImageJ). MFI of each islet was calculated by subtracting the mean intensity of a background region of interest (ROI). ImmTAAI co-localization was measured using the JaCOP plugin for ImageJ (*76*). Pearson's Coefficient was calculated between ImmTAAI and anti-ENTPD3 within a rectangular ROI comprising approximately 25% of the islet area, drawn to include labelled and unlabeled portions, while avoiding bright autofluorescent artifacts. T cell tracking was performed using the manual tracking mode for the TrackMate plugin for ImageJ (*77*).

**Generation of T cell clones.** T cell clones 4b and 15b specific for the pancreatic β-cell antigen $PPI_{6-14}$ (HLA-A*02:01 RLLPLLALL; 4b clone TCR has a KD ~800 µM, 15b clone TCR has a KD ~48 µM) were generated as described (*29*). T cell clone specificity was validated by dextramer staining (made in-house from biotinylated $PPI_{6-14}$ pHLA-A2 and fluorescently labelled Streptavidin-Dextramer®, Immundex ApS) and specific killing of EndoC-βH2 HLA-A*02 cells pulsed with $PPI_{6-14}$ peptide. The T cell clone 4b was transduced with lentivirus containing PDCD1 (PD-1) (OriGene #RC210364L1) as described (*29*). The clone was expanded with allogeneic irradiated PBMC from 3 donors in the presence of 1 µg/ml PHA-L and 100 U/ml IL-2 in T cell cloning media (RPMI 1640, 5% Human AB serum, 1% Penicillin/Streptomycin, 1% L-Glutamine, 1% MEM NEAA, and 1% Sodium Pyruvate).

**Flow cytometric assessment of avatar activation capacity.** The baseline activation status and capacity for T cell activation of the IGRP-reactive $CD8^+$ T cell avatars were established following 12 days of culture. Using approximately 1 x $10^6$ cells per condition, cells were stimulated with Dynabeads™ Human T-Expander CD3/CD28 at a 1:1 bead:cell ratio for either 0, 24, 48, or 72 hours. Following culture, avatars underwent viability staining with Live/Dead™ Near-IR viability dye (Thermo Fisher Scientific) for 10 minutes at 4˚C before washing with stain buffer (PBS + 2% FBS + 0.05% $NaN_3$ w/v). Before staining, cells were treated with TruStain FcX™ (BioLegend) for 5 minutes at 23˚C then stained with an extracellular antibody cocktail consisting of CD8-Alexa Fluor 700, CD69-BV421, and CD279-BV650 (clone and manufacturer information provided in Table S2) with Brilliant Stain Buffer Plus (Beckton Dickinson Biosciences) for 30 minutes at 4˚C. Data were collected on an Aurora 5L (16UV-16V-14B-10YG-8R) spectral flow cytometer (Cytek, Fremont, CA, USA), and analysis was conducted using FlowJo™ version 10.8.1 Software (BD Life Sciences).

**Human stem cell culture and sBC differentiation.** Undifferentiated human pluripotent stem (hPSC) $Mel1^{INS\text{-}GFP}$ reporter cells (*78*) were maintained on hESC qualified Cultrex (Biotechne #3434-005-002) in mTeSR+ media (STEMCELL Technologies #05826). Differentiation to stem cell-derived beta-like cells (sBC) was carried out in suspension-based, magnetic stirring platforms (Reprocell #ABBWVS03A-6, #ABBWVDW-1013, #ABBWBP03N0S-6) as described (*79, 80*). Briefly, 90% confluent hPSC cultures were dissociated into single-cell suspension by incubation with TrypLE (Gibco #12-604-021). Dissociation was halted with mTeSR+ media, and cells were counted using a Countess 3 cell counter (Thermo Fisher Scientific), followed by seeding 0.5 × $10^6$ cells/ml in mTeSR+ media supplemented with 10 µM ROCK inhibitor in spinner flasks. 3D sphere formation was performed for 48-72 hours. Differentiation media was changed daily by letting spheres settle by gravity for 3-5 minutes. Most supernatant was removed by aspiration; fresh media was added, and spinner flasks were placed back on stirrer system. sBC differentiation and cryopreservation were based on our published protocol (*81, 82*) with modifications as outlined below. Differentiation medias are as follows: induction of definitive endoderm differentiation using **d1 media** [RPMI containing 0.2% FBS, 1:5,000 ITS (Gibco #41400-045), 200 ng/ml Activin A (R&D Systems #338-AC-01M), and 3 µM CHIR99021 (STEMCELL Technologies #72054)] and **day 2-3**: RPMI containing 0.2% FBS, 1:2,000 ITS, and 100 ng/ml Activin A; **d4-5**: RPMI containing 2% FBS, 1:1,000 ITS, and 50 ng/ml KGF (Prepotech #100-19-1MG); **d6-7**: DMEM with 4.5 g/L D-glucose (Gibco #11960-044) containing 1:50 N-21 MAX (Biotechne #AR008), 1:100 NEAA (Gibco #11140-050), 1mM Sodium Pyruvate (Gibco #11360-070), 1:100 GlutaMAX (Gibco #35050-061), 3 nM TTNPB, (R&D Systems #0761), 250 nM Sant-1 (R&D Systems #1974), 250 nM LDN (STEMCELL Technologies #72149), 30 nM PMA (Sigma Aldrich #P1585-1MG), 50 µg/ml 2-phospho-L-ascorbic acid trisodium salt (VitC) (Sigma #49752-10G); **d8-9**: DMEM containing 1:50 N-21 MAX, 1:100 NEAA, 1 mM Sodium Pyruvate, 1:100 GlutaMAX, 100 ng/ml EGF (R&D Systems #236-EG-01M), 50 ng/ml KGF, and 50 µg/ml VitC; **d10-15**: DMEM containing 1:50 N-21 MAX, 1:100 NEAA, 1 mM Sodium Pyruvate, 1:100 GlutaMAX, 10 µg/ml Heparin (Sigma #H3149-250KU), 2 mM N-Acetyl-L-cysteine (Cysteine) (Sigma #A9165-25G), 10 µM Zinc sulfate heptahydrate (Zinc) (Sigma #Z0251-100g), 1x BME, 10 µM Alk5i II RepSox (R&D Systems #3742/50), 1 µM 3,3',5-Triiodo-L-thyronine sodium salt (T3) (Sigma #T6397), 0.5 µM LDN, 1 µM Gamma Secretase Inhibitor XX (XXi) (AsisChem #ASIS-0149) and 1:250 1 M NaOH to adjust pH to ~7.4; **d16-30**: CMRL (Gibco #11530-037) containing 1:50 N-21 MAX, 1:100 NEAA, 1:100 GlutaMAX, 10ug/ml Heparin, 2mM Cysteine, 10 µM Zinc, 1x BME, 1 µM T3, 50ug/ml VitC, 1:1000 Trace Elements A (Corning # 25-021-CI), 1:1000 Trace Elements B (Corning # 25-022-CI), 10 µM Alk5i II RepSox, and 1:250 NaOH to adjust pH to ~7.4. All media, except for mTeSR+, also contained 1x PenStrep.

**Cryopreservation and thawing of sBC.** Day 23 sBC were dissociated into single cells and cryopreserved as described (*53, 79*). Briefly, cells were quenched with 2% FBS in PBS and filtered using a cell strainer into FACS 5 mL tubes. Cells were counted using a Countess 3 cell counter (ThermoFisher Scientific) and resuspended at 3 x $10^6$ cells/100 µl of CryoStor® CS10 (STEMCELL Technologies). Cells were cryopreserved overnight before transferring to liquid nitrogen for long-term storage. For thawing, 1 mL of warm sBC media (d16-30 media described above) was added to the thawed cryovial dropwise before the entire volume was transferred into 5 mL of sBC media, counted, and seeded in Aggrewell 800 plates to generate clusters with 3,000 cells/cluster. After 24 hours, a partial media change was done to remove any debris. Fully formed clusters were generated after 48-72 hours.

**ImmTAAI sBC binding assay.** Day 23 sBC cryovials were thawed and plated into Aggrewell plates as described above with or without the presence of 20 nM ImmTAAI for 24 hours. Upon the partial media change at 24 hours, fresh ImmTAAI was added at 2 nM to the ImmTAAI treated cells or nothing as a control. After a total of 48 hours of reaggregation, half of the clusters were incubated for another 24 hours (total of 72 hours) with a re-dosing of 2 nM ImmTAAI while the other half were collected for imaging of ImmTAAI binding via confocal microscopy or dissociated into single cells and run live on the flow cytometer for quantification of ImmTAAI binding at the population level as detailed below. At 72 hours, these analyses were repeated with the additional clusters.

**sBC flow cytometry.** For sBC, single cell suspensions were made by washing clusters with PBS and incubating with 0.05% Trypsin with EDTA at 37°C for 12-15 minutes to create a single cell suspension. Cells were quenched with 2% FBS in PBS and filtered using a cell strainer into FACS 5ml tubes. Live cells were resuspended in FACS buffer for analyses on 5-laser Cytek Aurora for the pINS.GFP reporter marking beta cells and the ImmTAAI AF647. Analysis was performed using FloJo software v10.9 (BD Life Sciences).
